# A conserved mechanism for regulation of mtDNA copy number in eukaryotes

**DOI:** 10.64898/2026.05.29.727956

**Authors:** Flora McNulty, Andrew M. Shaw, Amy Shepherd, Jacqueline Tait-Mulder, Hannah Mearns, Ignacio Fernandez-Guerrero, Peggy Paschke, Elisabetta Tolla, Zaniah Gonzalez Galofre, Christina Wijo, Morgan McIntosh, Sergio Lilla, Graeme Clark, Alejandro Huerta Uribe, David Sumpton, Janneke Balk, Alberto Sanz Montero, Payam A. Gammage

## Abstract

Mitochondrial mass and mitochondrial DNA (mtDNA) copy number are coupled to metabolic demand at the cellular, tissue and organismal level, however, the molecular basis for homeostatic regulation of mtDNA is not understood. Here we show that mitochondria and mtDNA copy number are regulated by compartmentalisation of iron-sulfur (Fe-S) clusters, glutathione and cysteine, a mechanism we exemplify in model systems ranging from plants to human cells. Using genome-wide CRISPR screens we discovered that the mitochondrial ABC-family transporter, ABCB7, is a negative regulator of mtDNA copy number. Partial silencing of ABCB7 in human cells increased mtDNA 2-3 fold, enhancing mitochondrial mass and function. ABCB7 silencing compelled co-incident mitochondrial accumulation and cytosolic depletion of Fe-S clusters, simultaneously engaging the cellular iron starvation response and stabilising the mitochondrial glutathione transporter, SLC25A39. Transport of glutathione from the cytosol into mitochondria was co-incident with mitochondrial cysteine accumulation and cytosolic cysteine depletion, which was necessary and sufficient to increase mtDNA copy number in an integrated stress response-dependent fashion, with induction of PGC1β and ERRα. Silencing or partial loss of function mutations in the ABCB7 homologs of *D.melanogaster*, *S.cerevisiae* and *A. thaliana* elicited similar increases of mtDNA within these organisms. These data reveal a fundamental metabolic logic coupling compartmentalisation of redox co-factors to organellar genome content; a conserved axis across eukaryotes that pre-dates several elements of the mtDNA replication machinery.

## Main

Mitochondrial DNA (mtDNA) is maintained, replicated and expressed within the mammalian organelle by an orthologous complement of factors to those required for nuclear genome replication. Among numerous divergences between these two loci of eukaryotic genetics, how it is that stable, cell type-specific states of mtDNA copy number (mtCN) are maintained across several orders of magnitude to support the different metabolic requirements of tissues is not understood. Population-scale genomic analyses have revealed a complex interplay between mtCN and factors known to individually act as positive regulators of mtCN [1], and the overexpression of several factors required for mtDNA replication and transcription are known to induce altered states of mtCN [2–5]. Past attempts to discover factors regulating the initiation of mtDNA transcription and replication [1, 6–8] have robustly confirmed the role of positive regulators of mtCN. Canonical routes to stress-induced mitochondrial biogenesis downstream of AMPK signalling or environmental cues, such as PGC1α/ PGC1β-mediated activity of NRF1, form the current bedrock of understanding for how mtCN is controlled in response to stimulus, however these factors cannot fully explain homeostatic control of mtCN [9–14]. We therefore sought to identify novel homeostatic regulators of mtDNA using an unbiased screening approach.

Packaging of mammalian mtDNA into nucleoids with diverse states of accessibility and expression is an obligatory feature of mtDNA biology [15, 16]. The major protein component of the human mtDNA nucleoid is transcription factor A mitochondrial (TFAM), a protein with dual functionality that acts to initiate transcription of mtDNA and packaging of mtDNA [15–20] with approximately 1000 TFAM molecules bound per mtDNA molecule [2, 21]. As TFAM expression level is positively correlated with mtCN, we reasoned TFAM could be used as a reporter for mtDNA copy number in genome-scale, forward genetic screens. However, overexpression of TFAM is known to drive complex effects on mtDNA biology, with mild overexpression leading to increases in mtCN [22], and greater levels of overexpression known to result in compaction of mtDNA and loss of mitochondrial function [23]. To ensure physiological mtDNA regulation at the level of TFAM abundance, we engineered a TFAM reporter by modifying the endogenous locus of the gene. Using CRISPR/Cas9 nickase and a donor DNA construct, we engineered U2OS cells to bear 3xHA-Gly-linker-mCherry at the c-terminus of TFAM (**Figure 1a**). Selecting engineered cells by fluorescence-activated cell sorting (FACS) (**Figure 1b**), we isolated a homozygous knock-in clone for the reporter construct (hereafter referred to as KI/KI cells) (**Figure 1c,d**). The KI/KI cell line was validated through assessment of mitochondrial function and ethidium bromide depletion/repletion (**Extended Data Figure 1a-e**), which supported faithful TFAM function and localisation. To further validate the model for utility in pooled screening, we treated KI/KI cells with the known AMPK-driven mitochondrial biogenesis stimulator, AICAR, and also used CRISPR/Cas9 to knock out TFAM in KI/KI cells, observing expected directional fluorescence shifts by flow cytometry (**Extended Data Figure 1f,g**). Employing the Brunello CRISPR/Cas9 lentiviral library [24], we performed a genome-wide knockout screen in KI/KI cells, where the top and bottom 5% of fluorescent cells (hereafter TFAM-low and TFAM-high) were sorted using FACS following 13 days of puromycin selection, and were compared with the unsorted population for enrichment of sgRNAs associated with high or low TFAM expression (**Figure 1e**). Significant enrichment of sgRNAs for positive regulators of mtCN, such as essential components of the mtDNA maintenance, expression and replication machinery (*TFAM*, *MGME1* and *TFB2M*), in the TFAM-low population (**Supplementary Table 1**) technically validated the screening approach (**Figure 1f**), alongside which we observed enrichment of sgRNAs for *SLC25A28*, also known as mitoferrin 2, the mitochondrial Fe^2+^ importer in non-hematopoietic tissues. The gene with the most enriched sgRNAs in the TFAM-high population was *ABCB7*, a mitochondrial inner membrane-bound ABC-transporter [25, 26] (**Figure 1g**) (Supplementary Table 2).

**Figure 1.**
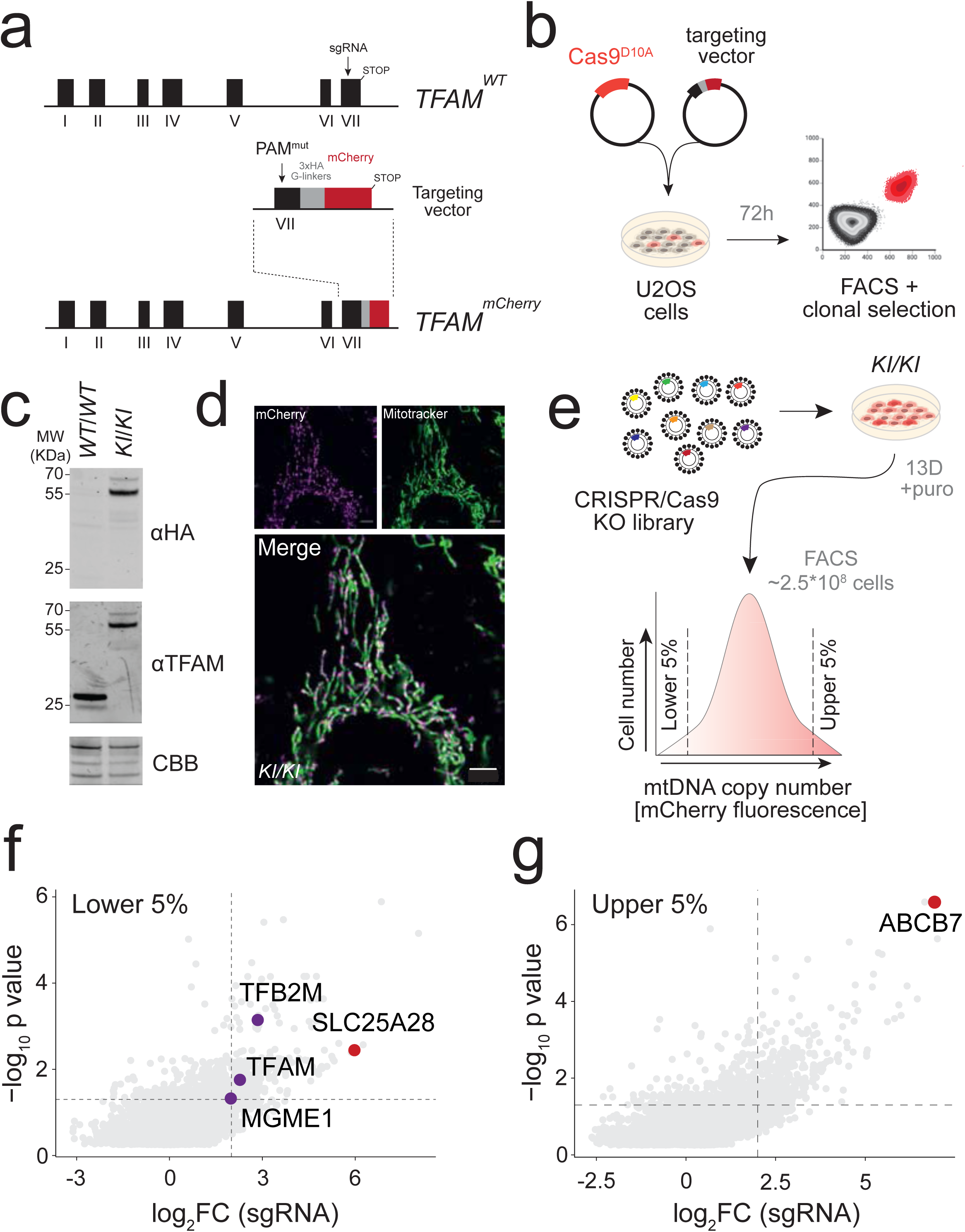
**A**. Map of human *TFAM* locus and recombination strategy. **B**. Workflow for generation of TFAM-mCherry knock-in U2OS cells. **C**. Immunoblotting of parental U2OS and isolated TFAM-mCherry KI/KI clone. CBB, Coomassie brilliant blue. **D**. Confocal microscopy of live KI/KI cells. Mitochondria visualised with MitoTracker green, native TFAM-mCherry signal detected. Scale bar, 3.4 μm. **E**. CRISPR screen strategy. Cells were infected with lentiviral particles and selected on puromycin for 13 days, then fixed and subjected to fluorescence-activated cell sorting (FACS) for mCherry fluorescence. **F**. Abundance of sgRNAs in lower 5% TFAM-mCherry fluorescence distribution. Thresholds for significance are log_2_ fold change < 2, adj-p value <0.05. **G**. Abundance of sgRNAs in upper 5% TFAM-mCherry fluorescence distribution. Thresholds for significance are log_2_ fold change < 2, adj-p value <0.05.

ABCB7 is structurally conserved from bacteria and throughout eukaryotes (**Figure 2a**), with the human transporter, yeast homolog, Atm1 (ABC transporter of mitochondria), and plant homolog, AtATM3, known to catalyse the ATP hydrolysis-linked transport of glutathione-conjugated Fe-S cluster intermediates (glutathione disulfide and glutathione-conjugated metals) unidirectionally from the mitochondrial matrix to the cytoplasm [25, 27–32]. In addition, partial loss-of-function mutations in ABCB7 are an established cause of ultra-rare X-linked sideroblastic anaemia with ataxia (XLSA/A), where significant non-heme iron deposits and mitochondrial proliferation are observed in the bone marrow by histopathology [26, 33–36], and missplicing of ABCB7 in SF3B1 mutant myelodysplastic syndrome similarly features non-heme iron-dense ringed sideroblasts [37]. Taken together these screen results implied that mitochondrial iron availability is essential for mtCN maintenance, and that retention of Fe-S clusters within mitochondria, or withholding Fe-S clusters from the cytosol, may dictate cellular mtCN. Fe-S cluster biosynthesis is thought to occur exclusively in mitochondria of non-photosynthetic eukaryotes, and Fe-S clusters are required by numerous proteins for diverse cellular functions, including DNA replication and repair (Polδ, Polɛ, primase), translational control (ABCE1) heme synthesis (ferrochelatase) and nucleotide synthesis (PPAT). In metazoans, cytosolic Fe-S clusters orchestrate cellular iron homeostasis through regulation of iron regulatory element (IRE) binding proteins IRP1 and IRP2, which bind IRE-containing mRNAs to increase labile iron. IRP1, in the Fe-S cluster absent apo-form, binds IRE-containing mRNAs to increase labile iron, while in the Fe-S integrated holo-state, IRP1 is otherwise known as ACO1, the cytosolic isoform of TCA cycle enzyme aconitase. IRP2 (also known as IREB2) stability is regulated by cytosolic Fe-S clusters via the SKP1-CUL1-F-box (SCF) ubiquitin ligase adaptor protein FBXL5, which requires an Fe-S cluster for this function [38–40]. As such, a feedforward loop, where iron starvation is sensed due to lack of cytosolic Fe-S clusters, resulting in uptake and delivery of iron to mitochondria to generate further Fe-S clusters, enabling mitochondrial biogenesis, provided an initial hypothetical framework to interrogate the function of ABCB7 in control of mtCN.

**Figure 2.**
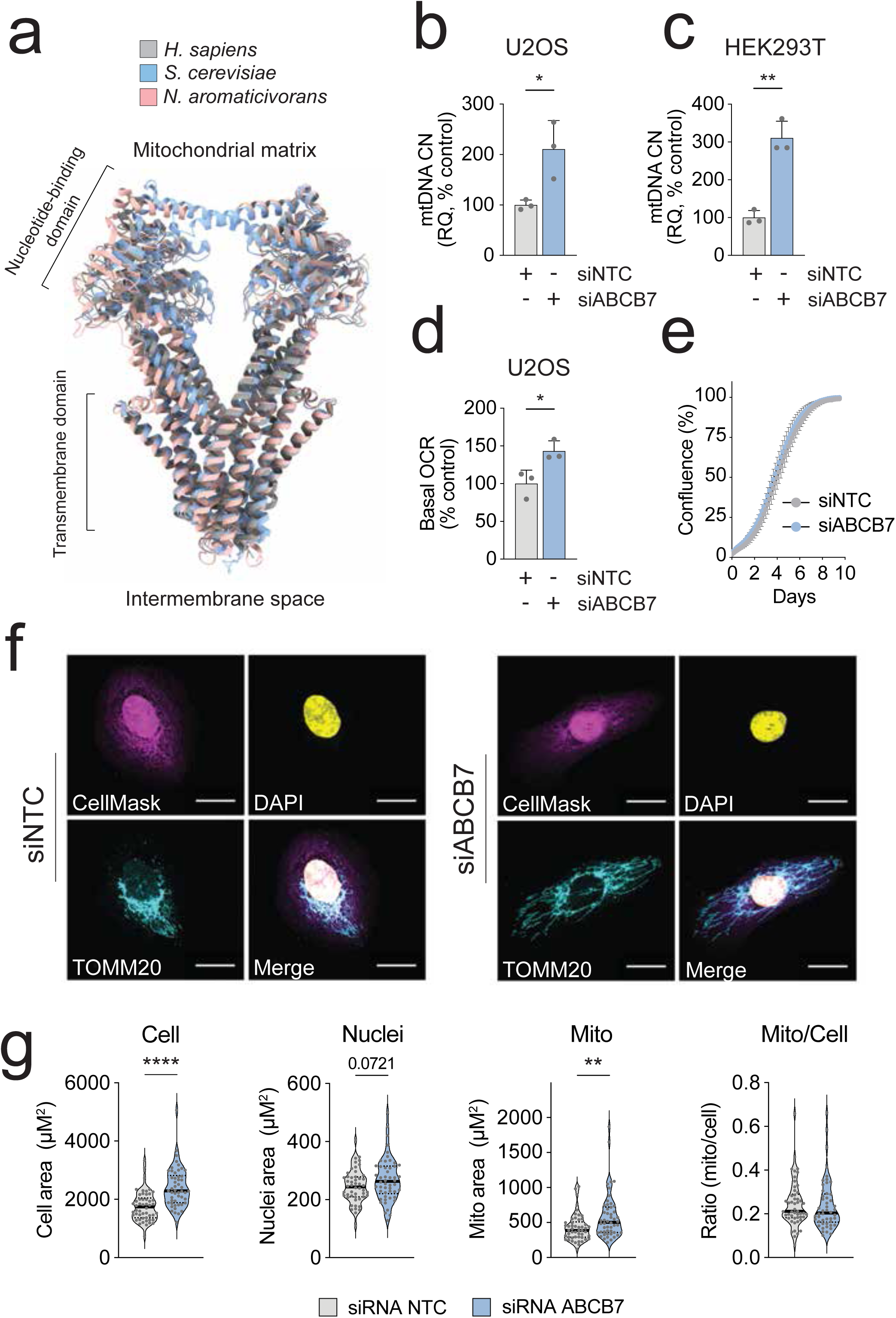
**A.** Structural comparison of human ABCB7 (gray) with yeast (blue) and bacterial (salmon) orthologs. **B.** Mean mtDNA copy number of U2OS cells treated with siRNA to *ABCB7* or non-targeting control (NTC) after 5 days. Each point represents the average mtDNA copy number of 3 separate culture wells from each of three independent experiments. Error bars, S.D. Welch’s t-test, * *p* <0.05. **C.** Mean mtDNA copy number of HEK293T cells treated with siRNA to *ABCB7* or NTC after 5 days. Each point represents the average mtDNA copy number of 3 separate culture wells from each of three independent experiments. Error bars, S.D. Welch’s t-test, * *p* <0.05. **D.** Mean oxygen consumption rate (OCR) measured in U2OS cells treated with siRNA to *ABCB7* or NTC after 5 days. Each point represents the average basal OCR across 12 separate wells from each of three independent experiments. Error bars, S.D. Welch’s t-test, * *p* <0.05. **E.** Cell proliferation measured with Incucyte. 12 separate wells measured per condition. Error bars, S.E.M. **F.** Example images from confocal microscopy of fixed U2OS cells treated with siRNA to *ABCB7* or NTC after 5 days. CellMask was used to define cellular boundary, DAPI to stain nuclei, TOMM20 to stain mitochondria. Scale bar 20μm. This experiment was repeated three times, a representative result is shown. **G.** Quantification of n=50 cells per condition. Violin plots indicate distribution, solid black line indicates mean, dashed line indicates interquartile range. Welch’s t-test. ** *p* < 0.01, **** *p* <0.0001.

Using publicly available data, we confirmed that *ABCB7* is an essential gene (**Extended Data Figure 2a-c**). Due to the essentiality of *ABCB7*, we experimentally validated our screen result by siRNA knockdown of ABCB7 in U2OS and HEK293T cells. A cell-type specific siRNA window was established, confirming that ABCB7 expression negatively regulates mtCN, with 2-3 fold increases observed in both cell lines after 5 days knockdown (**Figure 2b,c**) that demonstrated a non-linear relationship between extent of knockdown and changes in mtCN (**Extended Data Figure 2d,e**). Silencing of ABCB7 increased basal oxygen consumption rate (OCR) without impacting cell proliferation (**Figure 2d,e**), however an increase in cellular volume and trending increase in nuclear area were observed, with a proportional increase in mitochondrial mass (**Figure 2f,g**), consistent with previously described phenomena [41]. The proteome of ABCB7-depleted cells demonstrated clear divergence from control cells (**Extended Data Figure 3a**). Factors required for mtDNA replication and expression were generally upregulated at both the transcript and protein level (**Extended Data Figure 3b**) accompanied by transcriptional upregulation of PGC1β and ERRα, but not PGC1α or NRF1 (**Extended Data Figure 3c**). Complex effects on respiratory chain protein abundance were observed, with some mtDNA-encoded proteins being elevated (MT-ND5, MT-CO1) while others were depleted (MT-ATP8) (**Extended Data Figure 3d**). A number of nuclear encoded proteins of the respiratory chain were also depleted at the protein level, although this was not recapitulated at the level of gene expression (**Extended Data Figure 3e)**, suggesting post-translational regulation. Patterns of subunit depletion were not explained by their relative stage of incorporation during complex assembly, however trends in depletion of intermediate and early-stage assembly subunits were observed, possibly representing free pools of protein being exhausted in ABCB7 depleted cells (**Extended Data Figure 3f**). Taken together, these analyses suggest mitochondrial content and function within cells broadly scale with cell size, but do not scale in a linear fashion with increased mtCN following ABCB7 depletion.

Given the role of ABCB7 in providing cytosolic Fe-S clusters, the feedforward loop we predicted anticipated engagement of the iron starvation response in ABCB7 depletion (**Figure 3a**). Total intracellular labile iron was significantly increased in ABCB7-depleted cells (**Figure 3b**). Further, employing a cysteine oxidation state-sensitive proteomic workflow (**Figure 3c**) revealed depletion of ferritins (FTL, FTH1) and accumulation of IREB2 and transferrin receptor, TFRC, consistent with activation of the iron starvation response (**Figure 3d**) (**Supplementary Table 3**). Fe-S client proteins with lower abundance in ABCB7-depleted cells were almost exclusively located outside the mitochondrial matrix, including the essential DNA polymerase subunits POLD1 and POLE, DNA primase PRIM2, and cytosolic purine synthesis enzyme PPAT. Conversely, Fe-S proteins with increased abundance were exclusively located within the mitochondrial matrix, including Fe-S cluster assembly and chaperone components ISCU, FDX1 and CISD3. We further observed that key Fe-S ligand cysteines in POLD1, PRIM2 and PPAT were differentially reduced in ABCB7-depleted cells, contrasting with ligand cysteines in CISD3 and two subunits of Complex I, NDUFS2 and NDUFV1, which were differentially oxidised (**Figure 3e**), supporting a relative increase of Fe-S clusters within the mitochondrial matrix and depletion elsewhere in the cell.

**Figure 3.**
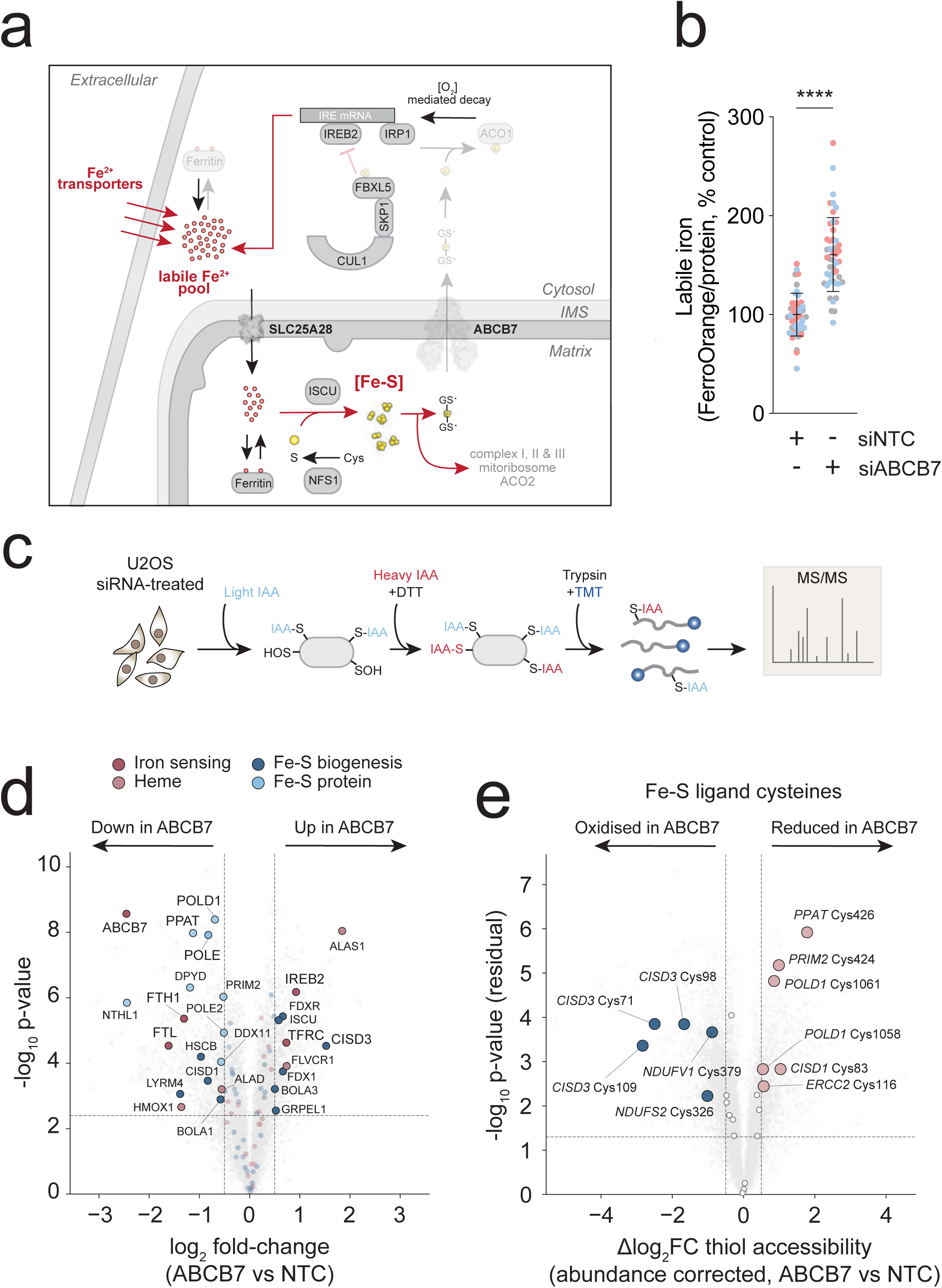
**A.** Proposed function of iron-sulfur (Fe-S) cluster export in cellular iron homeostasis and mitochondrial biogenesis. F-box protein, FBXL5, ubiquitinates IREB2 in an Fe-S-dependent manner. Aconitase apo vs holo states determine the role of this bifunctional protein as an iron-regulatory element-binding protein (apo, IRP1) or cytosolic TCA cycle enzyme, aconitase (holo, ACO1). A feed-forward loop driving Fe-S cluster accumulation in mitochondria is established by iron starvation responses being engaged, providing iron to mitochondria which do not release Fe-S clusters when ABCB7 is depleted, allowing the iron starvation response to persist. **B.** Mean labile iron measurement in U2OS cells treated with siRNA to *ABCB7* or non-targeting control (NTC) after 5 days, using FerroOrange probe. Normalised to total protein. Data points indicate measurements from 16 individual wells across three independent experiments. Colour indicates three independent experiments. Welch’s t-test, **** *p* < 0.001. **C.** Thiol redox-state sensitive proteomic workflow. Sequential treatment with light iodoacetamide (IAA) and heavy, isotopically labelled IAA + DTT labels thiols according to their native oxidation state. Tryptic peptides were labelled with isobaric tandem mass tags (TMT) and analysed by mass spectrometry (MS/MS). **D**. Differential iron-related protein abundances. Grey background, all detected proteins (n = 7,343). Coloured points, proteins from the iron-handling panel: two shades of red for the iron-sensing axis (IRP/IRE targets, regulators, iron transport, heme metabolism) and two shades of blue for Fe-S biology (biogenesis & trafficking, client enzymes). Significant hits (log₂FC > 0.5 and Perseus permutation FDR-corrected *p* < 0.05) are at full opacity with a black edge; non-significant panel proteins are dimmed. Significant gene symbols indicated. **E.** Fe-S ligand cysteine accessibility. Volcano plot of protein-abundance-corrected thiol accessibility. Grey background, all detected cysteines (n = 9,715). Filled coloured points, Fe-S cluster ligands. red = more reduced (NEM-accessible) in ABCB7 vs NTC blue = more oxidised (less NEM-accessible). Open circles, unchanged Fe-S ligand cysteines. *p*-values from Welch’s two-sample t-test on per-replicate residuals; horizontal dashed line, raw *p* = 0.05; vertical dashed lines, Δresidual significance <0.5 log_2_FC.

Fe-S cluster synthesis within mitochondria is a complex, multi-enzyme process, requiring several substrates and co-factors, including glutathione. Glutathione abundance in mitochondria has been shown to be regulated by Fe-S through the stabilisation of SLC25A39 [42–44]. In ABCB7-depleted cells, we observed that SLC25A39 was the most accumulated member of the SLC25 mitochondrial transporter family (**Figure 4a,b**) implying glutathione uptake from the cytosol into mitochondria may be required to maintain Fe-S synthesis. We confirmed mitochondrial accumulation of GSH and GSSG upon ABCB7 depletion by metabolite profiling of mitochondria isolated from 3xHA-mCherry-OMP25 expressing U2OS cells [45] (**Figure 4c,d**). To assess whether changes in mtCN were contingent upon SLC25A39, we generated SLC25A39 KO cell lines using CRISPR/Cas9 (**Extended Data Figure 4a**). Upon ABCB7 depletion, increased mtCN was not observed in SLC25A39KO cell lines, implying dependency (**Figure 4e**). However, overexpression of wild-type SLC25A39 or Δ72-86, a protease insensitive, constitutively active form of the carrier [43], were not sufficient to increase mtCN in wild-type or SLC25A39KO clone 3 (KO3) cells, establishing an epistatic relationship between ABCB7 and SLC25A39 (**Extended Data Figure 4b,c**). In addition, expression of the bacteria-derived, glutathione synthetase *GshF* in either mitochondria (mito*GshF*) or cytosol (cyto*GshF*) [42] did not induce changes in mtCN (**Extended Data Figure 4d,e**). Taken together, these data demonstrate that SLC25A39 stabilisation is necessary but not sufficient to increase mtCN.

**Figure 4.**
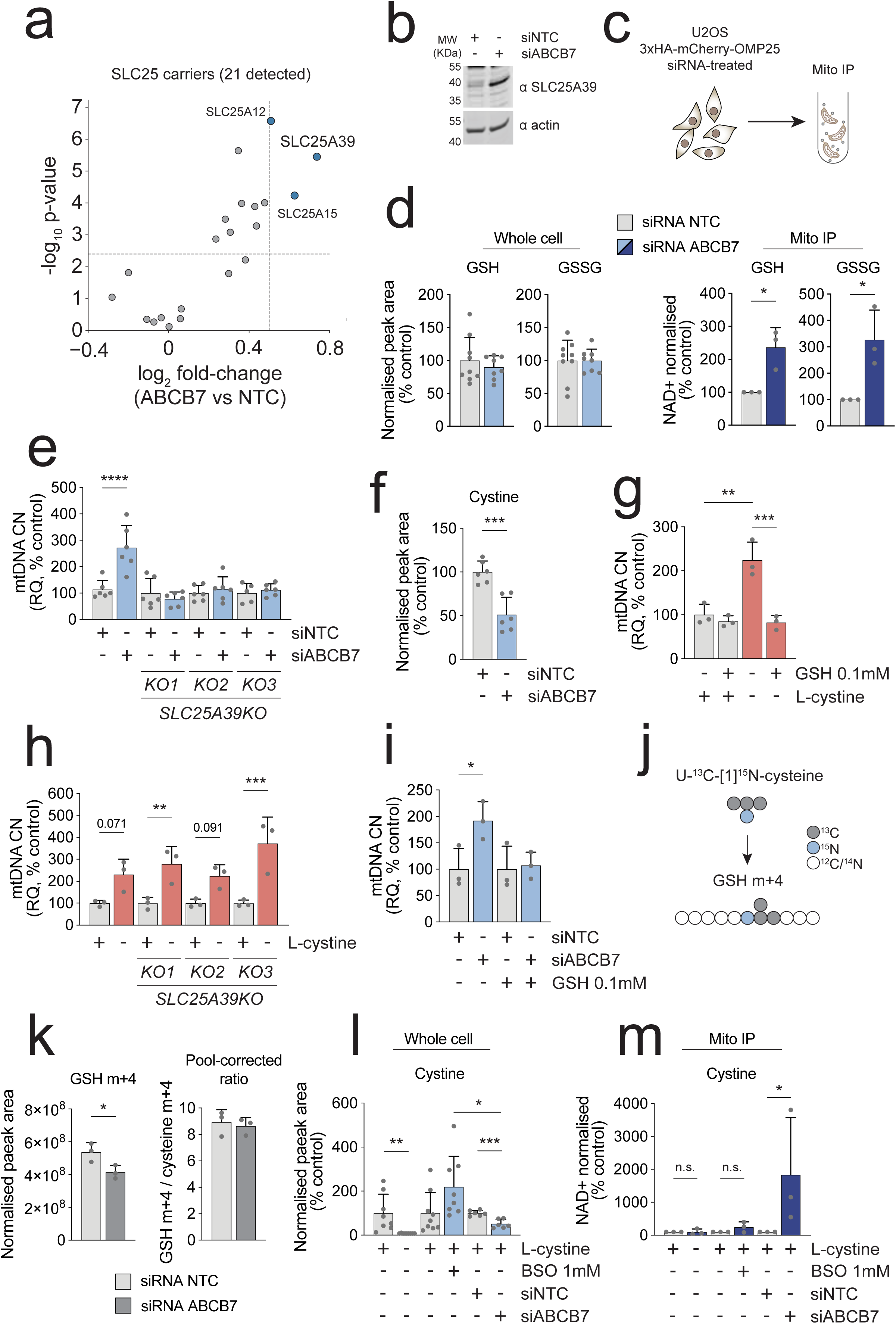
**A.** SLC25 family volcano plot restricted to detected SLC25 mitochondrial carrier family proteins (n = 21). Blue, significantly elevated. (log₂FC > 0.5 and Perseus FDR-corrected *p* < 0.05); grey, n.s. Italic gene labels on significant hits. **B.** Immunoblotting of U2OS cells treated with siRNA to *ABCB7* or non-targeting control (NTC) after 5 days. **C**. Scheme of mitochondrial immunoprecipitation (MitoIP) using U2OS cells stably expressing 3xHA-mCherry-OMP25. **D**. Mean abundance of reduced (GSH) and oxidised (GSSG) glutathione from whole cell (protein normalised) and MitoIP (NAD+ abundance normalised) metabolite extractions. Data points indicate three independent experiments. Error bars, S.D. Welch’s t-test, * *p* <0.05 **E.** mtDNA copy number of wild-type U2OS and *SLC25A39* knock out (KO) cells treated with siRNA to *ABCB7* or NTC after 5 days. Data points indicate mean of three wells from each of of six independent experiments. Error bars, S.D. Two-way ANOVA, Bonferroni corrected. **** *p* <0.0001. **F**. LC-MS determined cystine abundance in U2OS cells treated with siRNA to *ABCB7* or NTC after 5 days using NEM-spiked metabolomic extraction. Data points indicate individual wells across two independent experiments. Error bars, S.D. Welch’s t-test, *** *p* <0.01. **G**. Mean mtDNA copy number of wild-type U2OS cells grown in medium with or without L-cystine for 5 days, supplemented with 0.1mM GSH. Data points indicate mean of three wells from each of three independent experiments. Error bars, S.D. Two-way ANOVA, Bonferroni corrected. ** *p* <0.01, *** *p* < 0.001. **H**. Mean mtDNA copy number of wild-type U2OS and *SLC25A39KO* cells grown in medium with or without L-cystine for 5 days. Data points indicate average mtDNA copy number of 3 separate culture wells from each of three independent experiments. Error bars, S.D. Two-way ANOVA, Bonferroni corrected. * *p* < 0.05, ** *p* <0.01, *** *p* < 0.001 **I**. Mean mtDNA copy number of wild-type U2OS cells treated with siRNA to *ABCB7* or NTC after 5 days, supplemented with 0.1mM GSH. Data points indicate mean mtDNA copy number of three wells from each of three independent experiments. Error bars, S.D. Two-way ANOVA, Bonferroni corrected. * *p* < 0.05. **J**. Labelling fate diagram for U-^13^C-[1]^15^N-cystine. **K**. Mean protein normalised GSH m+4 and GSH m+4 : cysteine m+4 ratio. Data points indicate individual wells measured from a single experiment. Error bars, S.D. Welch’s t-test. * *p* < 0.05. **L**. Mean whole cell abundance of protein normalised cystine from U2OS cells grown without L-cystine, in the presence of BSO and treated with siRNA to ABCB7 or NTC after 5 days. Data points indicate individual wells from three (cystine, BSO) or two (siRNA) independent experiments. Error bars, S.D. Two-way ANOVA, Bonferroni corrected. * *p* < 0.05 **M**. NAD+ normalised mean of cystine abundance from mitochondria purified by mitoIP. Data points indicate three independent experiments. Error bars, S.D. Two-way ANOVA, Bonferroni corrected. * *p* < 0.05.

Metabolite profiling following ABCB7-depletion revealed that cellular NAD+:NADH, GSH:GSSG and energy charge were unchanged, however, the pyruvate:lactate ratio was diminished in ABCB7-depleted cells owing to a decrease in pyruvate abundance (**Extended Data Figure 4f-i**). Decreases of intracellular cystine (log_2_FC −0.47) and aspartate (log_2_FC - 0.75) with concomitant elevation of asparagine (log_2_FC 0.91), succinate (log_2_FC 1.11), acetyl-carnitine (log_2_FC 0.96) and choline (log_2_FC 0.96), among others, were also detected (**Extended Data Figure 4j**). These data suggested multiple electron donors could be accumulating due to an over-reduced ubiquinone (UQ) pool. Isotopic labelling with U-^13^C-glutamine indicated a modest increase in reverse flux of succinate dehydrogenase (SDH), derived from the ratio of succinate m+3 : m+4 relative to the ratio of aKG m+3 : m+5 (**Extended Data Figure 5a,b**). Multimodal profiling of ABCB7-depleted cells further revealed a shift towards lipid synthesis and storage over oxidation (**Extended Data Figure 5c**) with notable increase in abundance of PPARG transcripts **(Extended Data Figure 3c)**. Taken together, these data suggest an electron-saturated UQ pool present in ABCB7-depleted cells, evidenced by increased succinate abundance, reverse SDH flux and likely diminished electron entry via ETFDH, stalling β-oxidation to underpin acetyl-carnitine accumulation.

Gene set enrichment analysis (GSEA) of ABCB7-depleted cells revealed robust increases of integrated stress response (ISR), MYC and mTORC1 signalling, underpinned by significant depletion of DEPTOR, LAMTOR2 and TFEB with increased MYC and MAX expression observed at both RNA and protein level (**Extended Data Figure 6a,b**) (**Supplementary Table 4**). Consistent with the absence of AMPK pathway activation, AMPK component abundance was either unchanged or demonstrated modest, non-coordinated regulation, with no change in AMPK phosphorylation detected (**Extended Data 6c,d**). A limited form of ISR signalling, characterised by elevated ATF3/4/5, ASNS, CHAC1, SLC7A11 and SLC3A2 was also observed in ABCB7-depleted cells (**Extended Data Figure 6e,f**), with canonical UPR markers, including HSPA5, HSP90B1 and ATF6 targets remaining unchanged, suggesting absence of broader ER-stress. GCN2 (EIF2AK4) was the sole ISR effector kinase observed to be significantly elevated at transcriptional and protein level, with HRI (EIF2AK1) concomitantly repressed (**Extended Data Figure 6f**). Increases in mtCN following depletion of ABCB7 were prevented by treatment of cells with ISRIB (**Extended Data Figure 6g**), blocking eIF2a-mediated inactivation of eIF2B. In addition, an established negative regulator of ISR signalling at the level of translational control, BZW1 [46], was a highly significant hit in the TFAM-high population of the initial CRISPR screen identifying ABCB7 (**Extended Data Figure 6h**). Together, these data implied that ISR signalling, likely via amino acid stress through GCN2, is a key component of the ABCB7-depleted mtCN-elevated phenotype.

Depletion of intracellular cystine coupled to upregulated expression of the glutamate-cystine exchanger, ^x^CT (SLC7A11), its obligatory heavy chain partner, SLC3A2, the cytosolic glutathione catabolising enzyme, CHAC1, and a GCN2-driven ISR suggested cytosolic cysteine-depletion may serve as a signal linking accumulation of mitochondrial Fe-S and SLC25A39-mediated uptake of GSH with ISR activation. We further confirmed depletion of cystine upon silencing of ABCB7 using a workflow incorporating NEM-treatment to preserve endogenous thiol oxidation states (**Figure 4f**), and then tested the causality of this effect by culturing cells in L-cystine-free medium, observing a ∼2 fold increase in mtDNA over 5 days that was rescued by supplementing cells with 0.1mM GSH (**Figure 4g**). A precipitous decline in GSH and GSSG was observed in L-cystine starved cells (**Extended Data Figure 7a,b**), however, cells treated with buthionine sulfoximine (BSO), inhibiting GSH synthesis, demonstrated an equivalent decline in GSH (GSSG undetectable) without increased mtCN (**Extended Data Figure 7c,d**). Notably, SLC25A39 abundance was elevated in both BSO and L-cystine starvation (**Extended Data Figure 4a, Extended Data Figure 7e**), however SLC25A39 KO cells cultured in medium without L-cystine still demonstrated a robust increase of mtCN (**Figure 4h**), strongly supporting the notion that cystine depletion is a downstream effector. This finding was further supported by observation of comparable trends in mtCN when cells are treated with the ^x^CT inhibitor, erastin (**Extended Data Figure 7F**). In keeping with these results, supplementation of ABCB7-depleted cells with 0.1mM GSH prevented increases in mtCN (**Figure 4i**), underpinning the importance of cysteine depletion-mediated ISR signalling in the regulation of mtCN.

While an increased proportion of cellular GSH and GSSG were located within mitochondria following ABCB7 depletion, total cellular abundance of GSH and GSSG was not impacted (**Figure 4d**). Cysteine/cystine are maintained at several orders of magnitude lower concentration within cells relative to glutathione precursors glycine and glutamate [47]. Thus, we reasoned compensatory cytosolic glutathione synthesis, owing to disproportionate glutathione compartmentalisation within mitochondria, and increased expression of CHAC1, might lead to proportionally greater depletions of cystine in the cytosol without meaningfully impacting glutamate, glycine or total glutathione pool size. To test this possibility cells were cultured in medium with isotopically labelled U-^13^C-[1]^15^N-cystine, where we observed a decrease in GSH m+4 labelling in ABCB7-depleted cells, with the cysteine m+4 pool size-adjusted fraction of GSH m+4 unchanged between NTC and ABCB7 depleted cells (**Figure 4j,k**). To better understand compartmentalisation effects, we measured metabolites from purified mitochondria of L-cystine starved cells, cells supplemented with BSO and cells treated with siRNA to *ABCB7*, revealing an ∼18-fold increase in mitochondrial cystine only in conditions of ABCB7 depletion (**Figure 4l,m**). This accumulation was GSH-dependent, as mitochondria purified from BSO-treated cells do not accumulate cystine despite increased abundance of cystine in BSO-treated cells. Previously it has been observed that CHAC1 is partially localised to mitochondria, where in conditions of cysteine starvation it acts to maintain the mitochondrial cysteine pool, required for Fe-S cluster biogenesis via NFS1, consuming GSH in the process [48]. In keeping with this, a substantial increase in the mitochondrial abundance of alanine, the product of cysteine desulfuration by NFS1, was also detected in cysteine starved and ABCB7 depleted conditions (**Extended Data Figure 7G**). Taken together these data robustly support ABCB7 depletion-induced, SLC25A39-dependent mitochondrial compartmentalisation of GSH and cystine, which simultaneously act as a signal to increase mtCN in the cytosol via the ISR while supporting mitochondrial Fe-S biogenesis.

ABCB7 and SLC25A39 are deeply conserved across eukaryotes (**Extended Data Figure 8a**) (**Supplementary Table 5**). To examine the broader evolutionary relevance of the ABCB7/SLC25A39 axis, we examined conservation of pathway function using genetic models of the fruit fly *Drosophila melanogaster*, Baker’s yeast *Saccharomyces cerevisiae* and the model plant *Arabidopsis thaliana* (**Figure 5a**). We tested the effect of ubiquitous (da-Gal4) RNAi knockdown of *ABCB7* in *D. melanogaster* larvae, where ∼70% depletion of the mRNA (**Extended Data Figure 8b**) resulted in a ∼2-fold increase of mtCN at larval stage L3 across the entire organism (**Fig. 5b,c**), with subsequent failure to pupate. Using *S. cerevisiae* cells where the endogenous promoter of the *ABCB7* homolog, *ATM1*, has been replaced with an inducible *GAL1-10* promoter, growth on medium containing glucose (YPD), which represses promoter activity, resulted in a ∼1.5 fold increase in mtCN (**Figure 5d,e**). Finally, we assessed the impact of a partial loss of function mutant, *atm3-1*, that bears a deletion in the nucleotide-binding domain (ΔNBD), of the *A. thaliana ABCB7* homolog, *ATM3/ABCB25* [28]. Assessing total seedling DNA from *atm3-1* plants compared to wild types we observed ∼2.5 fold increases in mtCN across four genes of *A. thaliana* mtDNA (**Figure 5f,g**).

**Figure 5.**
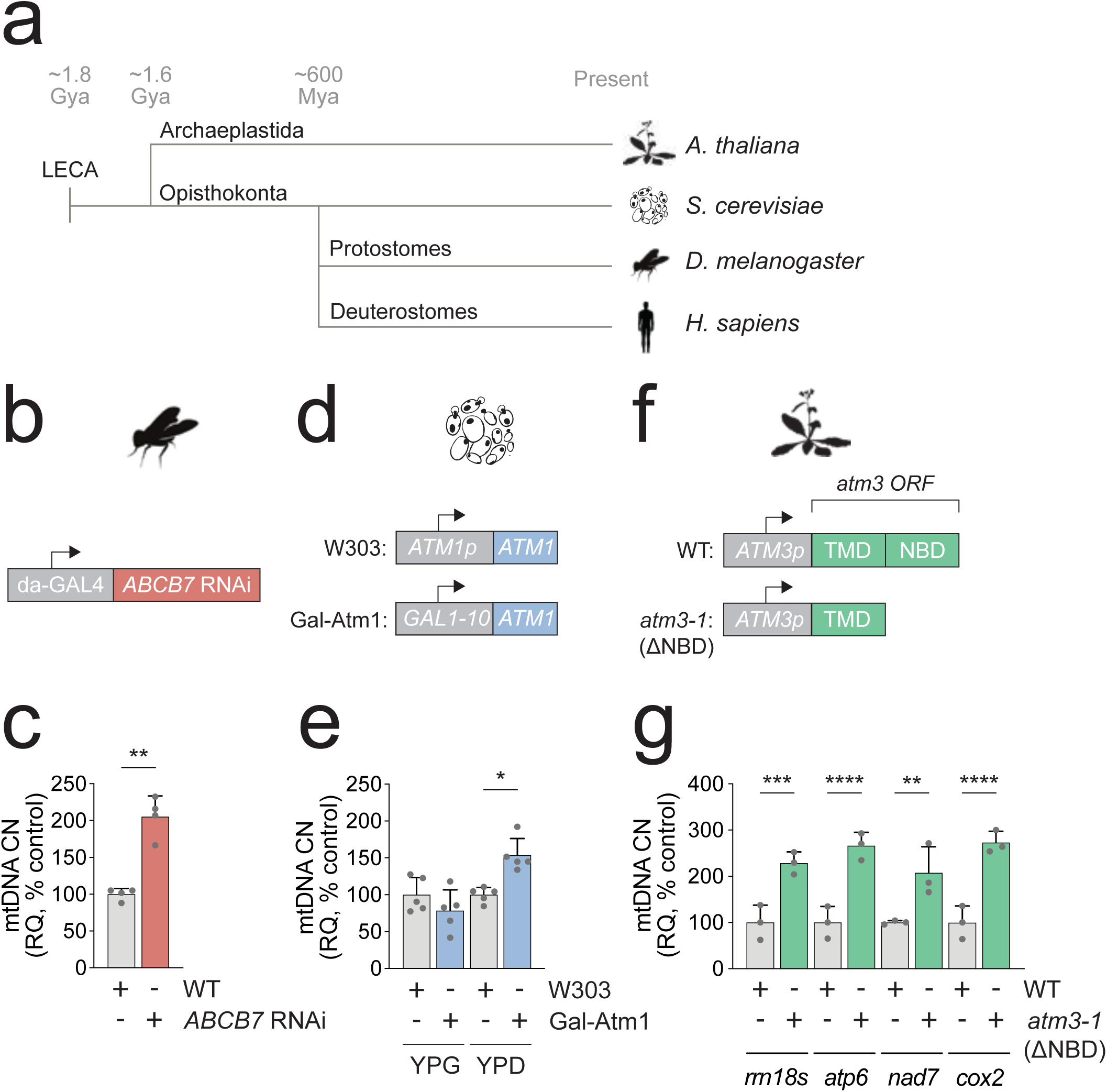
**A.** Phylogenetic diagram indicating evolutionary distance and divergence of model systems. LECA, last eukaryotic common ancestor. **B.** Schematic of transgenic *D. melanogaster* da-Gal4 *ABCB7* RNAi allele. **C.** Mean mtDNA copy number of L3 larvae. Data points indicate individual larvae (n=4). Error bars, S.D. Welch’s t-test. ** *p* < 0.01. **D.** Schematic of *S. cerevisiae* Atm1p and Gal1-10 promoter alleles. **E**. Mean mtDNA copy number of five independent streakings of strains grown on indicated carbon source. Data points indicate average of three measurements from each of five independent streakings. Error bars, S.D. Two-way ANOVA, Bonferroni corrected. * *p* < 0.05. **F.** Schematic of *A. thaliana atm3* (WT) and *atm3-1* alleles, indicating transmembrane domain (TMD) and nucleotide binding domain (NBD). **G.** Mean mtDNA copy number of three independently cultivated seedling cohorts. Four seedlings per cohort, including roots and leaves, were combined per independent cohort, and each of these was measured three times. Data points indicate average mtDNA copy number per seedling cohort across four distinct regions of *A. thaliana* mtDNA. Error bars, S.D. Welch’s t-test.** *p* < 0.01, *** *p* < 0.001, **** *p* < 0.0001.

## Discussion

The regulation of mitochondrial mass and mtDNA copy number is fundamental to eukaryotic life, and a longstanding area of interest in basic and translational research. It is established that acute sensing of energetic crisis or environmental challenge can drive mitochondrial biogenesis through AMPK and PGC1α/NRF1 [11–14]. Factors necessary for the replication and expression of mtDNA in animals, including TFAM, POLG, and the mitochondrial replicative helicase Twinkle, have been extensively characterised, with their manipulation known to alter mtCN [2–5]. However, we believe these factors represent the effector machinery of mtDNA replication rather than its regulatory logic. Moreover, the maintenance and expression apparatus for mtDNA are among the most divergent features of mitochondrial biology across eukaryotes; a pattern that is difficult to reconcile with a model by which mtCN is principally governed at the level of the replisome. Past efforts have also focused on understanding potential mitochondrial retrograde signalling molecules, such as reactive oxygen species (ROS), as influencers of mitochondrial biogenesis [49]. Here, we report a previously unrecognised dimension through which mitochondrial content and metabolic states are regulated across all major eukaryotic lineages tested: compartmentalisation of glutathione, cystine and Fe-S clusters.

The extent of GSH-dependent mitochondrial compartmentalisation of cystine upon depletion of ABCB7 was a stark and unanticipated result. Considering theoretical mass balance and relative compartment volumes, these data suggest that the majority of cellular cystine may be located within mitochondria in conditions of ABCB7 depletion. Cystine accumulation within mitochondria was seen to be dependent upon GSH synthesis, and GSH has previously been observed to buffer mitochondrial cysteine levels, enabling Fe-S cluster synthesis during cysteine starvation through the activity of CHAC1 [48]. It therefore appears that depletion of cysteine within the cytosol and concurrent accumulation of cysteine within mitochondria serve dual purposes; underpinning Fe-S synthesis whilst simultaneously driving ISR-dependent signalling and transcriptional programmes to enhance mitochondrial biogenesis (**Figure 6a,b**). That ABCB7 depletion simultaneously engages the iron starvation response, ensuring accumulation of both essential substrates required for Fe-S biogenesis in mitochondria, iron and cysteine, argues for an Fe-S-centric view of mitochondrial biogenesis. How such disequilibrium between cytosolic and mitochondrial cysteine can be maintained remains an open question, and the lack of compensatory GSH synthesis underpinning cystine depletion in ABCB7 silenced cells suggests that depletion of cellular cystine may be the result of a more complex series of events. It is possible that ABCB7 itself represents a major route of mitochondrial GSH efflux, rebalancing the mitochondrial and cytosolic cysteine pools. It may also be the case that a lack of detectable NRF2-driven programs to increase cellular cystine uptake underpins cellular cystine depletion in these states.

**Figure 6.**
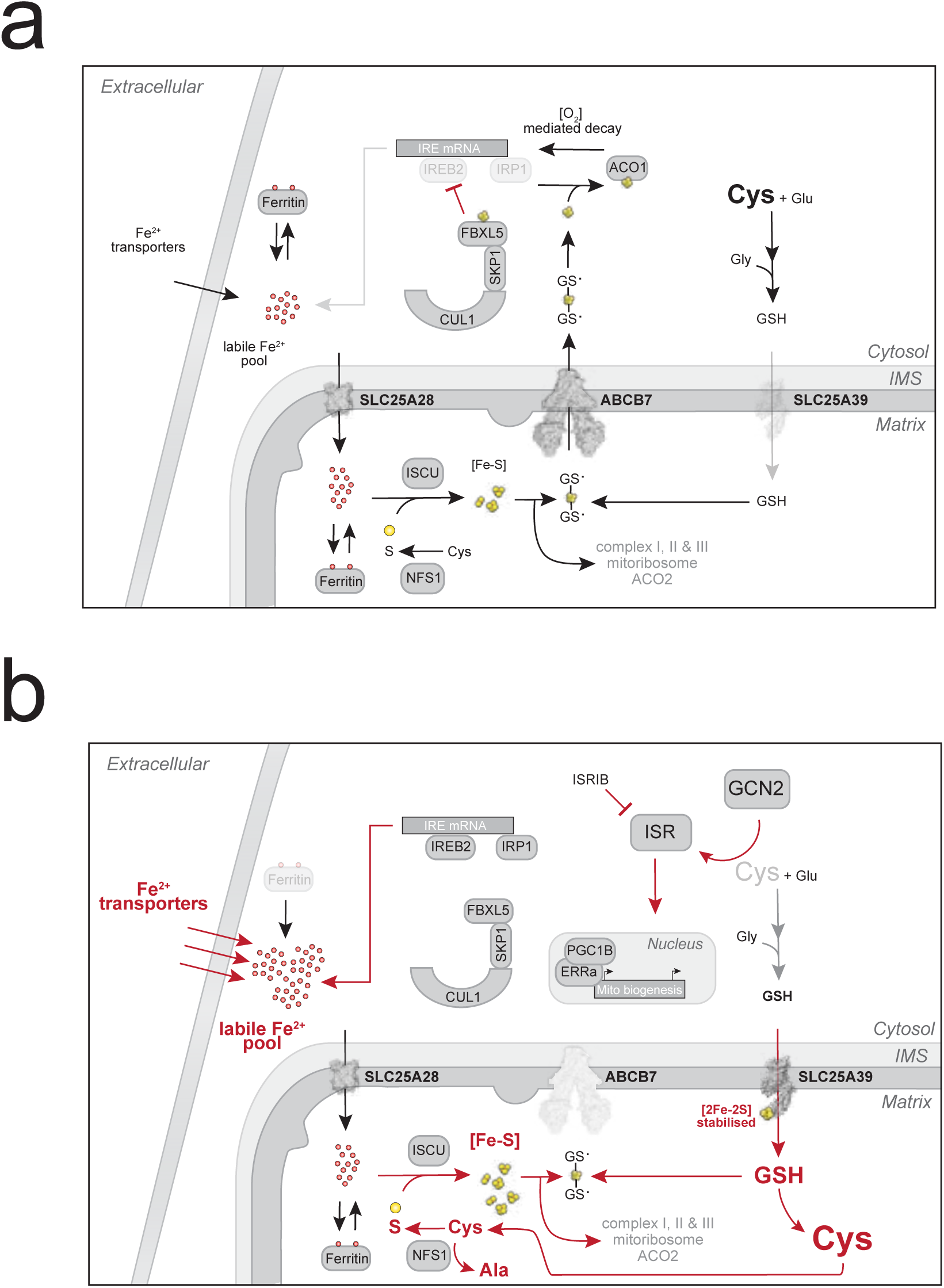
**A.** Schematic of ABCB7 mediating iron homeostasis through Fe-S cluster export, resulting in concomitantly reduced IREB2 and IRP1 levels and reduced SLC25A39 stability. **B.** Schematic of the mechanistic basis for sensing and elevation of mitochondrial and mtDNA copy number resulting from decreased ABCB7 expression. Iron starvation response is activated due to decreased efflux of Fe-S clusters from mitochondria, resulting in stabilisation of IREB2, IRP1 and SLC25A39. GSH and cystine accumulate in mitochondria, supplying sulfur for Fe-S biogenesis, accompanied by cytosolic cystine depletion that is sensed by integrated stress response (ISR) effector kinase, GCN2, to drive cellular adaptations in an ISRIB-sensitive fashion, resulting in mitochondrial biogenesis via PGC1β and ERRα.

A striking feature of the biogenesis programme downstream of ABCB7 is that this appears to operate via ISR induction and through PGC-1β/ERRα rather than the better established PGC-1α/NRF1 axis (**Extended Data Figure 3c**). PGC-1α is canonically induced under conditions of acute metabolic stress, such as cold exposure, exercise, and energetic deficit. In contrast, PGC-1β/ERRα engagement, possibly downstream of mTORC1 and MYC, positions this programme within the constitutive nutrient-sensing and growth-control circuitry of the cell, consistent with a homeostatic mechanism that tunes mitochondrial mass in proportion to ongoing metabolic demand. This scaling appears to arise primarily through metabolic fluxes — the demand for Fe-S clusters determining their own supply — rather than through a transcriptional programme directed at the replication or biogenesis apparatus. In this manner, mitochondrial function is set not only by how much mtDNA replication machinery is expressed, but by the extent of metabolic activity the organelle is called upon to support.

The capacity to maintain redox homeostasis is an essential feature of life, although diversity in the mechanistic basis for how this is achieved is readily observed throughout eukaryotes. Iron, for example, in both the elemental state and the form of Fe-S clusters, is tightly regulated through functionally analogous but mechanistically diverse processes between plants and metazoans [50]. Given the diversity of eukaryotes, and their respective mitochondrial genetic systems, it has been presumed that regulation of cellular mitochondrial content might operate under similarly divergent mechanisms. For example, mtDNA in humans is a genetically compact, circular molecule of 16.5kb that does not undergo meaningful recombination or transpositional events, encoding membrane-bound OXPHOS protein subunits. However, the variation within mitochondrial genetic systems across eukaryotes is vast, including organisms with megabase genomes, linear genomes, sub-genomic fragments and frequent rearrangements from which a diversity of soluble, monomeric and insoluble, OXPHOS complex-associated proteins are expressed [51]. Subsequently, the factors controlling mitochondrial genome replication, expression and maintenance are highly divergent, with several proteins controlling these processes in humans, including PGC1-α/β, the mtDNA replicative polymerase γ and TFAM, emerging only in opisthokonta and bilateria (**Extended Data Figure 7b**). The data we report here strongly support an alternate view: mitochondrial content is not regulated solely by the replisome machinery, but rather through the ABCB7-SLC25A39 axis, a principle of cell biology that has been under selection since LECA.

Considering the profound diversity in mitochondrial genome structure and the factors required for their regulation, repair and expression throughout eukaryotes, it is parsimonious that the regulatory logic and core apparatus governing mitochondrial genomes would arise prior to the specific machinery required for mtDNA replication or expression in any given organism. Owing to the comparably ancient origins and functional interplay of these systems, it may seem intuitive that mTOR, ISR, iron and redox homeostasis should prominently feature in the biogenesis programme of mitochondria, although the apparent decoupling of Fe-S based regulation of iron homeostasis from mitochondrial biogenesis in plants [28] suggests a degree of flexibility in this dimension. Our data suggest that coupling of mitochondrial biogenesis and availability of labile iron is perhaps a later evolutionary elaboration in opisthokonts, providing mechanistic understanding of the pathogenic mechanism by which partial loss of function mutations in ABCB7 and missplicing in myelodysplastic syndrome result in pathological iron accumulation in the haematopoietic system [37].

While ABCB7 has been under strong positive selection throughout eukaryotic evolution, positioned at the apex of Fe-S biogenesis and mediating cellular supply of Fe-S cluster intermediates, it is not clear that ABCB7 itself represents the precise physiological locus of mtCN regulation. The rate of Fe-S transport through ABCB7 theoretically integrates mitochondrial ATP:ADP:AMP, iron availability, cytosolic and mitochondrial cysteine abundance, mitochondrial glutathione abundance, mitochondrial Fe-S demand/supply, and likely through O_2_-dependent Fe-S decay, oxygen sensing. Our data further suggest that ubiquinone pool size and redox state are key constraints on mitochondrial function, regardless of mtCN. It is therefore plausible that convergence of these inputs, their dependencies (rate limiting processes within each input) and further unidentified contingencies determine mtDNA copy number across tissues and organisms, with ABCB7 serving as an essential node through which these signals are relayed, rather than the sole point at which the circuit is regulated. However, this previously undiscovered link between mitochondrial metabolism, glutathione, cysteine and iron biology provides a new lens through which diverse metabolic states, from stem cells to immune cells, and even cancer cells, might be viewed.

In summary, we present evidence for a conserved regulatory mechanism underpinning mitochondrial biogenesis and the control of mitochondrial genome copy number that is responsive to nutrient availability and metabolic states, forming an ancient basis for the maintenance of mitochondrial homeostasis across eukaryotes

## Methods

### Cell culture

U2OS WT (ATCC) KI/KI and HEK293T (ATCC) cells were maintained in Dulbecco’s Modified eagle medium (DMEM) supplemented with GlutaMAX (Gibco, 10566016), 20% Fetal Bovine Serum (Gibco, 10270-106), 1% Penicillin/Streptomycin (Gibco, 15070063), 0.1% uridine (Merck, U3003-50g). Cells were grown at 37°C at 5% CO2 and split as required. All cells were authenticated by in house authentication service through morphology and STR profiling. All cells were tested repeatedly for mycoplasma, and were negative.

### Media and drug treatments

For experiments, U2OS or HEK293T cells were seeded at a low density. The following day medium was substituted with regular growth medium or for cystine deprived conditions with high glucose DMEM, lacking glutamine, methionine, or cystine (Gibco, 21023024) supplemented with 20% dialysed FBS (Gibco, 26400044), 100 µg m^−1^ uridine (Merck, U3003-50g), 0.11 g l^−1^ sodium pyruvate (Gibco, 11360070), 2 mM L-glutamine (Gibco, 25030081) and 1% Penicillin/Streptomycin (Gibco, 15070063), 0.2mM L-Methionine (Sigma Aldrich, M5308). Cells were treated as indicated in figures (and legends) with the following chemicals: BSO (Sigma Aldrich, B2515), GSH (Sigma Aldrich G4251), erastin (MedChemExpress, HY15763), ethidium bromide (Life Technologies, 15585-01), ISRIB (MedChemExpress, 548470-11-7). Cells were incubated at 37°C and 5% CO2.

### Cell line engineering

#### Generation of TFAM-mCherry KI U2OS cells

Oligos were designed to target the terminal exon of TFAM. They were annealed at 95°C for 5 minutes and then a gradual reduction (1 degrees/min) back to room temperature. These were cloned into pSaGuide, a gift from Kiran Musunuru (Addgene plasmid # 64710; http://n2t.net/addgene:64710; RRID:Addgene_64710). Plasmids expressing CAG-SaCas9-WPRE, a gift from Timo Otonkoski (Addgene plasmid # 89996; http://n2t.net/addgene:89996; RRID:Addgene_89996). U2OS cells were transfected with *TFAM* exon 7 sgRNA and the donor template were transfected into U2OS cells at a 3:1:1 donor:sgRNA:Cas9 ratio using Lipofectamine 3000 (Invitrogen, L3000001). 72 hours after transfection cells were sorted for mCherry expression. Three days after, cells were trypsinised, counted and single cells clones generated using limiting dilution in 96 well plates using sterile filtered conditioned medium (50/50 fresh/conditioned medium) from U2OS wild type cells. Single cell clones were then screened via confocal microscopy to select promising clones. Selected clones were assessed further by confocal microscopy using a Zeiss LSM880 AiryscanFast Microscope, western blotting and flow cytometry.

#### SLC25A39 knockout U2OS KI/KI cells

sgRNA SLC25A39 sequences (see table) were designed using [52] annealed and cloned into pSpCas9(BB)-2A-Puro (PX459) V2.0 (Addgene Plasmid #62988) using the BbsI cloning site (New England Biolabs, #3539). U2OS TFAM KI/KI cells were transfected with the empty pSpCas9(BB)-2A-Puro (PX459) plasmid or the one containing the SLC25A39 sgRNA and selected with puromycin. After selection, cells were counted and single cell clones generated using limiting dilution in 96 well plates using sterile filtered conditioned medium (50/50 fresh/conditioned medium) from WT U2OS cells. Single cell clones were then screened through RNA isolation and RT-qPCR for SLC25A39 expression. Three SLC25A39 clones and one empty vector clone were selected for further analysis.

#### SLC25A39KO TFAM KI/KI cells expressing SCL25A39 wild-type or Δ72-86 mutant

HEK293FT cells were transfected with FLAG-SLC25A39wt-pLix401-Blast, FLAG-SLC25A39Δ72-86-pLix401-Blast (Genewiz) were transfected into HEK293FT using Lipofectamine 2000. Viral supernatant was filter sterilised and added to SLC25A39KO3 and the control clone cells for 48 hours after addition of 6μg/mL polybrene (Merck, TR-1003). Cells with integrated plasmid were selected using 10ug/ml blasticidin (Life Technologies) and bulk populations were characterised by western blot after doxycycline induced expression of SLC25A39wt or SLC25A39Δ72-86.

#### Mito-tag U2OS cell generation

Mito-tag U2OS cells were generated using virus produced in Phoenix-ampho cells following transfection with pMXs-IRES-Bla-3HA-EGFP-OMP25 (David Sabatini, Addgene Plasmid #83356) using Lipofectamine 2000 according to the manufacturer’s instructions and as above. Cells with integrated plasmid were selected using 10ug/ml blasticidin and checked for GFP and HA expression by western blot, flow cytometry and fluorescence microscopy.

Oligo table:

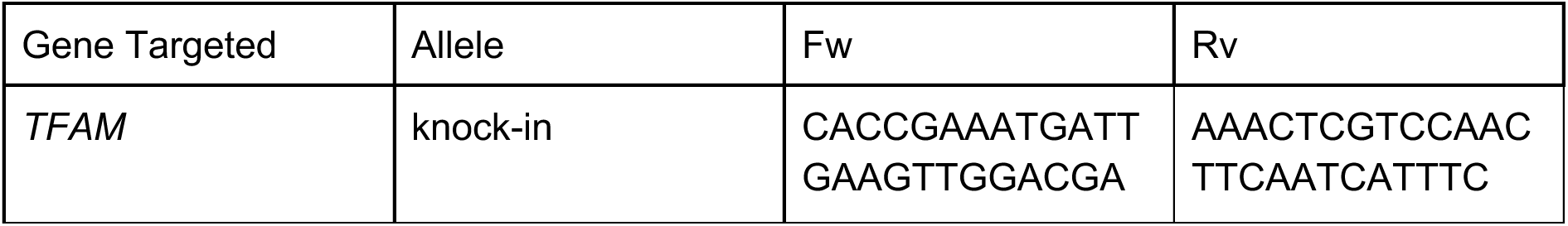
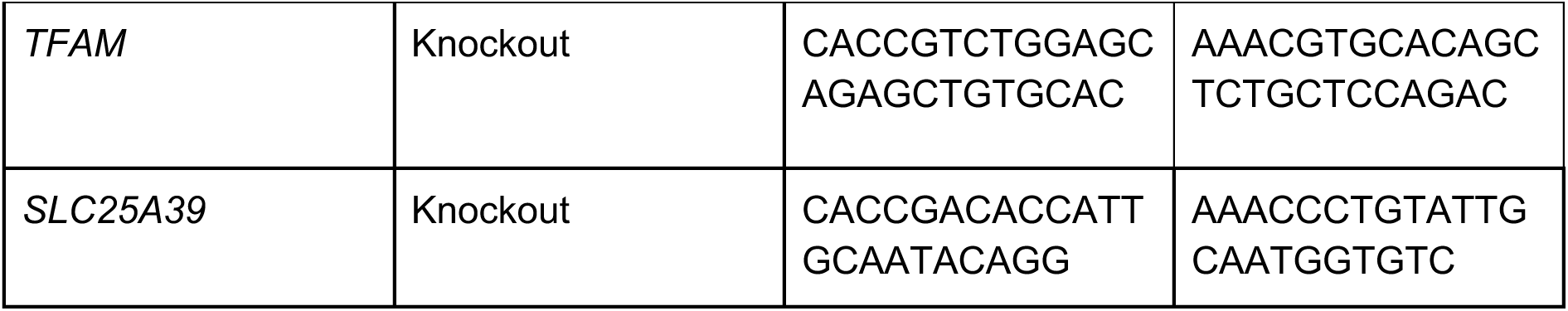

### Immunoblotting

Western blots were performed as previously published [53]. In some cases, protein concentration per sample was determined by DC Protein Assay (Bio-Rad Laboratories) in other cases using a BCA protein assay (Life Technologies). Equal amounts of proteins were loaded onto precast SDS–PAGE 4–12% Bis-Tris Bolt gels (Life Technologies) after which proteins were transferred to nitrocellulose using a Mini Trans-Bolt Cell (Bio-Rad Laboratories). Nitrocellulose membranes were stained with Ponceau S solution to visualise equal loading and an image was taken. After destaining using TBS-T, membranes were blocked in 5% milk (Marvel, Original Dried Skimmed Milk) in TBS-T for 1 hour and then incubated overnight with indicated primary antibodies: TFAM (Protein Tech, 22586-1-AP), HA (Roche, 11867423001), OXPHOS Human WB antibody cocktail (Abcam, ab110411), SLC25A39 (Protein Tech, 14963-1-ap), FLAG (Sigma, F3165), TOMM20 (Abcam, ab186735), Vinculin (Abcam, ab109235), beta-actin (Abcam, ab8226), AMPK (Cell Signalling Technologies, #2532) or phospho-AMPK (Cell Signalling Technologies, #2531). Primary antibodies were diluted in milk or in the case of phospho antibodies in BSA, both in TBS-T. The following day membranes were briefly washed using TBS-T and incubated with secondary antibodies: Donkey anti-Mouse 800CW (LI-COR Biosciences, 926-32212), Donkey anti-Mouse 680RD (LI-COR Biosciences, 926-68072), Donkey anti-Rabbit 800CW (LI-COR Biosciences, 926-332213), Donkey anti-rabbit 680RD (LI-COR Biosciences, 926-68073), Goat anti-Rat 800CW (LI-COR Biosciences, 926-32219) or Goat anti-Rat 680RD (LI-COR Biosciences, 926-68076) at 1:5000 dilution in TBS-T for 1 hour. After washing with TBS-T, membranes were analysed using an Odyssey CLx imager and Image Studio Lite software (v.6 LI-COR Biosciences).

### Flow cytometry

Prior to flow cytometry, KI/KI U2OS cells were seeded and either transfected or treated as described in the figure legends. Cells were transfected at a ratio of 5μg DNA:7.5μL using Lipofectamine 3000 (Life Technologies, L3000001). Samples were stained with DAPI stain 1:1000 (Life Technologies, 62248) and run on a BD LSRFortessa X20 (BD Biosciences) instrument. mCherry excitation, 561nm laser with 600LP and 610/20 BP filters; DAPI excitation, 405nm laser with a 450/50 BP filter. Data were analysed using FlowJo (v.10.10.1).

### CRISPR KO screen

KI/KI U2OS cells were seeded at ∼50% confluence in 5-layer cell culture flasks (Scientific Laboratory supplies, 353144) with 20mL of cell culture medium per layer. Cells were transduced at an MOI of 0.3 with 6μg/mL polybrene (Merck, TR-1003) using the Brunello Library, a gift from David Root and John Doench (Addgene, #73178). After 48 hours, the viral medium was removed and replaced with cell culture medium supplemented with 2μg/mL puromycin (Gibco, A1113803). Cells were split as required. After 14 days, cells were fixed at room temperature in the dark using a 30-minute incubation, in accordance with the eBioscience™ Intracellular Fixation & Permeabilization Buffer Set (Thermo Fisher Scientific, 00-5523-00). Fixed samples were either sorted or cells were kept as an unsorted control. Samples were stored at 4°C in PBS supplemented with 2% FBS (Gibco, A5256701) until they were sorted using FACS. Sorting gates were set to collect the top 5% and bottom 5% of fluorescent mCherry cells. Cells were sorted using a BD FACSAria III (BD Biosciences) instrument with mCherry excited using 561nm laser with 600LP and 610/20 BP filters.

Extraction from fixed cells proceeded from adapted from [54] and using the QIAamp DNA mini kit (QIAGEN, 51306). Phusion High-Fidelity PCR kit (Invitrogen, F553L) was used to amplify sgRNAs. Reactions contained 30 ng of DNA, 1x Phusion buffer, 200 μM dNTPs, 0.5 μM forward primer, 0.5 μM of reverse primer, 5 mM MgCl_2_, 0.02 U/μL Phusion polymerase. Primers contained partial Nextera adaptors for subsequent library preparation. The PCR program used the following conditions: initial denaturation 98°C for 30 seconds, followed by 35 cycles of denaturation at 98°C for 10 seconds, annealing temperature at 59°C for 30 seconds, and extension time of 10 seconds. A final extension of 72°C for 7 minutes was performed. PCR products were purified using 1.8X beads in accordance with the AMPure Protocol (Beckman, A63881).

Following first round PCR, full length libraries were generated using Nextera XT CD Indexes (Illumina, 20018708) and amplified using repliQa HiFi ToughMix PCR mastermix (Invitrogen). Resultant libraries were then purified using 1X AMPure magnetic beads (Beckman Coulter). Post library QC was then performed using High Sensitivity D1000 screentape (Agilent) for library sizing and profiling and quantified using Qubit High Sensitivity DNA assay. Libraries were then pooled equimolar, to a final concentration of 2nM prior to sequencing. The library pool was then sequenced on an Illumina NextSeq 2000 P4 100 cycle Flow-cell, at Single Read 75bp read length. FastQ files were generated on-board using Dragen BCLConvert (Illumina).

Raw single-end sequencing reads were trimmed using Cutadapt (v.4.9) [55] to extract the 20bp spacer sequences. The sequencing reads from multiple PCR reactions obtained for the same samples were catenated. FastQC (v.0.12.1) [56] and MultiQC (v.1.17)[57] were used to assess read quality. sgRNA counts and ranked significance was determined using MAGeCK (v.0.5.9.5) [58]. Control-based normalisation using the 1000 non-targeting control guide RNAs in the Brunello library was employed. Sufficient library representation was achieved with >94% of guides identified and a GINI-index of 0.185 in unsorted control. Differences in gene abundances were determined by comparing sorted populations with unsorted control, using the RRA algorithm. Output was filtered based on detection of 2 or more sgRNAs per gene, with duplicates enriched at both ends of the screen removed as technical artefacts. Filtered lists of enriched sgRNAs in the low and high population are available in the supplementary information (Supplementary Table 1, Supplementary Table 2).

### siRNA knockdown

Knockdown of between 3-5% of wild-type mRNA level was required for ABCB7 phenotypes. Each batch of siRNA was re-optimised for transfection before experimental use. Cells were seeded at 50,000 cells in a 6-well plate for transfection the next day. For transfection, Lipofectamine RNAiMAX Transfection Reagent (Invitrogen, 13778150) was diluted in Opti-MEM (Gibco, 31985062). The siRNA, either NTC (non-targeting control) siRNA (ON-TARGETplus Control pool, Non-Targeting pool, Dharmacon, D-00180-10-05) or *ABCB7* siRNA (ON-TARGETplus SMART pool, Human *ABCB7*, Dharmacon, L-007305-00-0010), was first diluted to 10 μM in water as a stock and then for transfection diluted in Opti-MEM to working concentration. The diluted siRNA was then added to the diluted RNAiMAX, gently mixed and incubated for 5 minutes at room temperature. The siRNA mixture was then added dropwise to the cells and left for the time indicated in the figure legends.

### Protein structure alignment

Experimentally determined structures of ABCB7 and orthologs were accessed from PDB (Accession numbers: 7VGF, 4MYC, 6VQU). Structures were processed using ChimeraX to remove nucleotide-binding domain ligands and pseudobond annotations. MatchMaker, using default settings, was applied to align the structures using *H.sapiens* ABCB7 as the reference structure.

### DepMap analysis

Cell-line CRISPR essentiality and gene expression data were obtained from the Cancer Dependency Map (depmap.org) [59, 60]. *ABCB7* CRISPR gene effect scores (Chronos) and dependency probabilities were extracted for all profiled cell lines (n = 1,208). Lines were classified as dependent at a probability threshold of 0.5 and as common-essential at a Chronos gene effect of ≤ −1. Bulk RNA-seq expression values (log₂(TPM+1) were filtered to one entry per model using the IsDefaultEntryForModel flag (n = 1,719 models).

### mtDNA copy number analysis

DNA was extracted from each sample (DNeasy Blood & Tissue Kit, Qiagen). DNA extractions were measured in duplicate, or single DNA samples in triplicate, as indicated in the figure legends. Droplets were generated in 20 μl reactions containing 1 ng of isolated DNA, 100 nM of indicated forward and reverse primers, 10 µl of QX200 ddPCR EvaGreen Supermix (Bio-Rad Laboratories) and water. Reaction droplets were then subjected to PCR and analysed using the QX200 Droplet Digital PCR System (Bio-Rad Laboratories). For human cells nuclear DNA primers against *NCOA3* a ofnd mitochondrial DNA primers against *MT-TL1* and a primer annealing temperature of 60 °C was used. For *Saccharomyces cerevisiae* samples, nuclear primers amplified *COX1* and mitochondrial primers amplified *CDC28* with an annealing temperature of 65°C. For *Arabidopsis thaliana* samples, nuclear primers amplified *AtAct1* and mitochondrial primers amplified *AtRrn18s, AtAtp6, AtNad7* and *AtCox2* using an annealing temperature of 60°C.

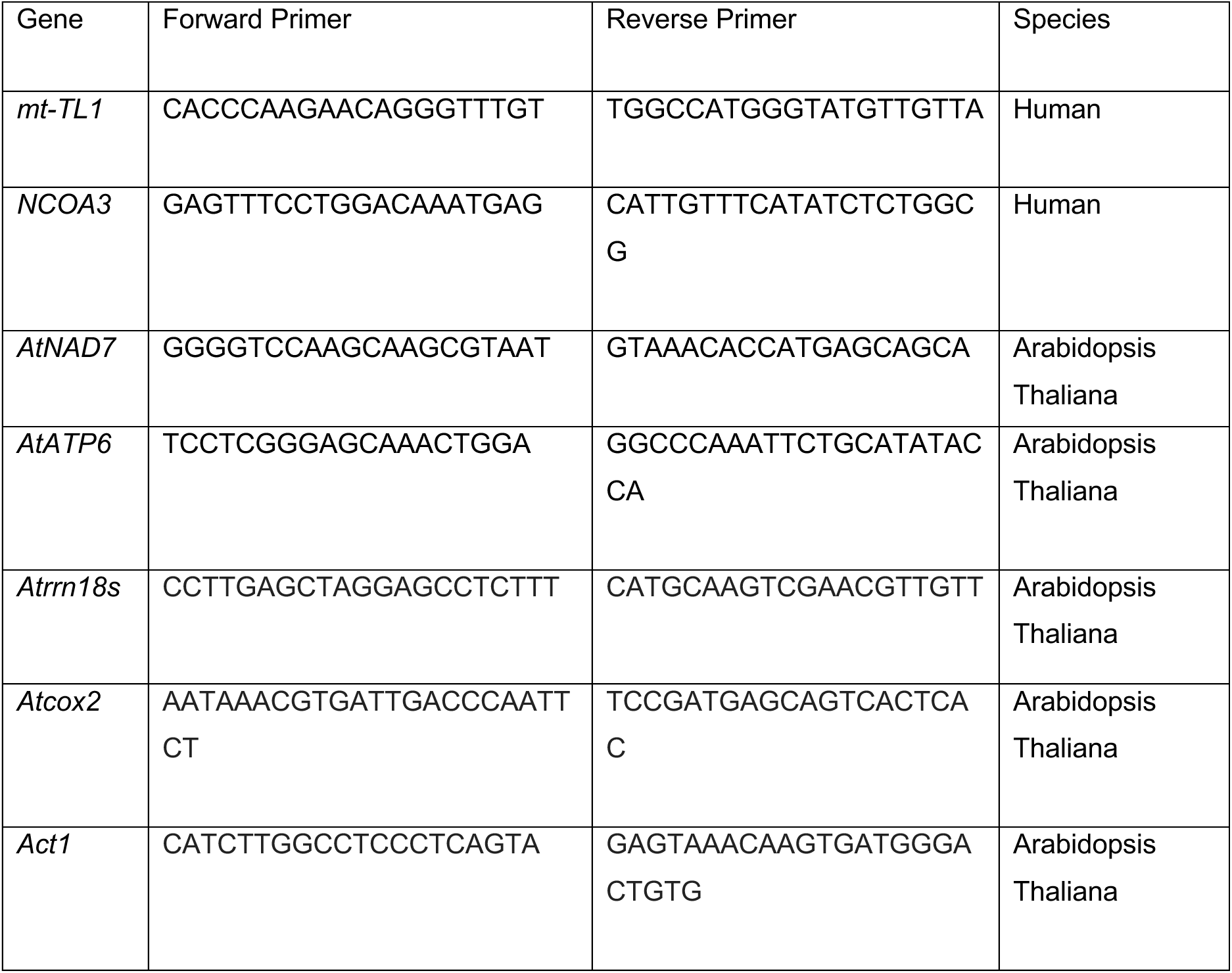
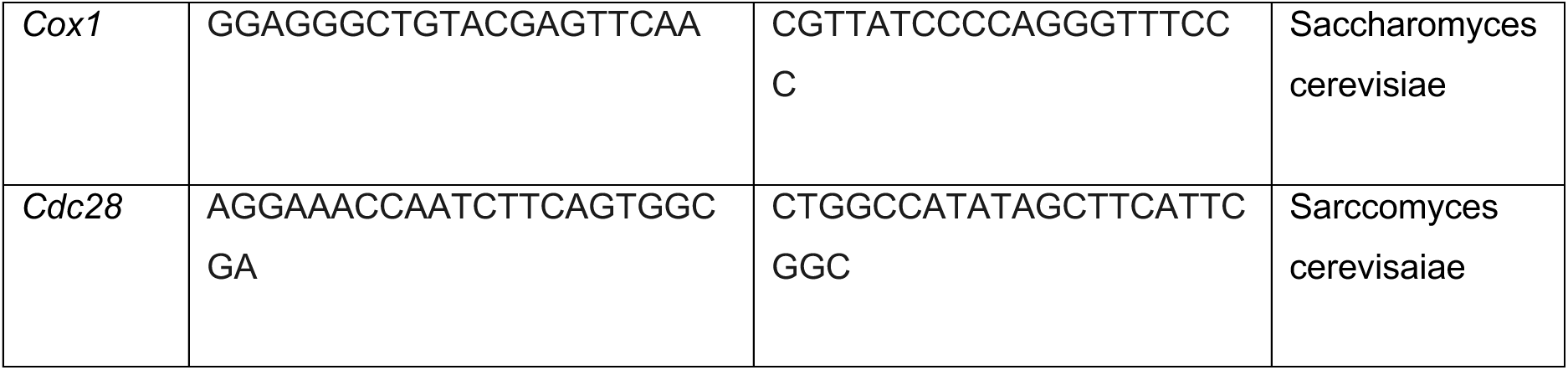

*MT-TL1* and *NCOA*3 primers are from [61]. *mt-Cox1* and *Cdc28* primers are from [62]. *AtNad7,AtAtp6* and *AtAct1* primers are from [63]. *Atrrn18s* and *AtCox2* primers are from [64].

#### mtDNA copy number - D. melanogaster

MtDNA copy number was quantified by qPCR using total DNA extracted by phenol–chloroform. Relative mtDNA copy number was determined by amplifying the mitochondrial *mt:lrRNA* (16S rRNA) gene (forward: 5′-TGGCCGCAGTATTTTGACTG-3′; reverse: 5′-TCGTCCAACCATTCATTCCA-3′) and normalising to the nuclear reference gene *RpL32* (forward: 5′-AGGCCCAAGATCGTGAAGAA-3′; reverse: 5′-TGTGCACCAGGAACTTCTTGAA-3′). Reactions were performed using FAST SYBR Green Master Mix on a QuantStudio 3 Real-Time PCR System under standard cycling and melt-curve conditions. Each reaction was performed in technical triplicate, with four independent biological replicates per group. Ct values were averaged and analysed using the ΔΔCt method. Statistical significance was assessed using a two-tailed Student’s t-test.

### Reverse transcription-quantitative PCR (RT-qPCR)

RNA was extracted using the RNeasy Mini Kit (QIAGEN, 74104) and QIAshredder columns (QIAGEN, 79654). RNA was converted to cDNA using the High-Capacity RNA-to-cDNA Kit (Applied Biosystems, 4387406). The 20 μl reverse transcription reaction consisted of: 10 μl of 2x RT Buffer mix, 1 μl of 20x RT enzyme mix, up to 9 μl of RNA sample and then nuclease-free water to bring the total to 20 μl. The thermocycler (ProFlex PCR system, Applied Biosystems) conditions were: 37**°**C for 1 hour, 95**°**C for 5 minutes and then held at 4**°**C until analysis. cDNA was diluted in nuclease-free water as required. Triplicate samples were analysed in duplicate. Each reaction contained 12.5 μl 1x Taqman Universal PCR Master Mix (Applied Biosystems, 4364340) with 1.25 μL of Taqman probes (included forward and reverse), in nuclease-free water in a total volume of 25 μl. 20 μl was loaded per well in the qPCR plate, with 3 μl of cDNA added per well. Taqman probes targeted *ABCB7* (Hs00188776) and *B2M* (Hs00968432). The qPCR (Quant Studio 3, Applied Biosystems) conditions under the standard run mode, cover set to 105**°**C, were: up 1.6**°**C/s to 50**°**C (for 2 minutes), up 1.6**°**C/s to 95**°**C (for 10 minutes), then 40 cycles of up 1.6**°**C/s to 95**°**C (for 15 seconds) and down 1.6**°**C/s to 60**°**C (for 1 minute). *B2M* was used as the reference gene and the 2^(-ΔΔCt) method was used to calculate the relative expression of *ABCB7*.

#### RT-qPCR - D.melanogaster

*ABCB7* mRNA levels were quantified by qPCR using cDNA synthesised from total RNA extracted with TRIzol reagent (Thermo Fisher Scientific) according to the manufacturer’s instructions. Complementary DNA (cDNA) was generated using the High-Capacity cDNA Reverse Transcription Kit and used as template for qPCR. Relative *ABCB7* expression was determined by amplifying *ABCB7* (forward: 5′-CTAAGGGACCGGGAACACAC-3′; reverse: 5′-TGGCATCGCTGCTGTTCTTA-3′) and normalising to *RpL32* (forward: 5′-AGGCCCAAGATCGTGAAGAA-3′; reverse: 5′-TGTGCACCAGGAACTTCTTGAA-3′).

Reactions were performed using FAST SYBR Green Master Mix on a QuantStudio 3 Real-Time PCR System under standard cycling and melt-curve conditions. Each reaction was performed in technical triplicate, with four independent biological replicates per group. Ct values were averaged and analysed using the standard curve method. Statistical significance was assessed using a two-tailed Student’s t-test.

### Oxygen consumption analysis

The basal OCR measurements were performed using a Seahorse XF Cell Mito Stress Test (Agilent) as per the manufacturer’s instructions. In brief, after 4 days, siRNA transfected or treated cells were plated at 25.000 cells per well in a XFe96 plate (Agilent). The following day, at day 5, cells were incubated for 1 h at 37 °C in bicarbonate-free 200 µl Seahorse XF Medium with the addition of 1% FBS, 25 mM glucose, 1 mM sodium pyruvate and 2 mM glutamine. The OCR was determined in triplicate measurements every 9 minutes using a XFe96 Analyzer (Seahorse Bioscience) at baseline and after addition of each drug in the following order and concentration: oligomycin (1 μM), CCCP (1 μM), rotenone (1 μM) and antimycin A (1 μM) (all from Sigma). Upon completion, cells were washed with 200ul PBS, aspirated and the plate was put in −80C and a protein Bradford assay (Biorad, #5000006) was performed later to normalise the fluorescence reading protein concentrations of each well.

### Cell proliferation assays

U2OS cells were transfected with siRNA. The following day they were reseeded at a density of 500 cells/well in a 96-well cell culture dish in 200 uL of cell culture medium. Incucyte images were collected on an Incucyte S3 (Sartorius), using a 10x/0.3 Plan Fluor objective lens (Nikon) and brightfield illumination. Images were collected with an image size of 1024 × 1024 pixels, yielding a pixel size of 1.24 x 1.24 um. 5 positions per well were captured every four hours. The instrument was located inside an incubator (HERAcell 240i, ThermoFisher Scientific) maintained at 37 degrees and 5% CO2. Images were acquired using the Incucyte software (Sartorius). Images were analysed using Incucyte software (v. 2025B), using the Basic Analyzer algorithm. The Segmentation algorithm was chosen as “AI Confluence”, with a minimum area filter of 1120 um^2 and a maximum eccentricity of 0.98.

### Confocal microscopy

U2OS cells transfected in a 6-well cell culture dish, were reseeded in fibronectin (Sigma-Aldrich, F1141) coated glass bottomed 6-well cull culture dishes 4 days after transfection at a densities of 10 000, 20 000 and 40 000 cells per well. The following day, cells were fixed in 4% formaldehyde for 10 minutes. Cells were permeabilised through incubation with 20mM glycine (Sigma-Aldrich, G7126), 0.05% Tritonx100 (Sigma-Aldrich, X100-5mL) for 10 minutes. Cells were blocked in 4% BSA (Sigma-Aldrich, A7030) in PBS. Fixed cells were incubated overnight with anti-TOMM20 (Abcam, ab186735) at 4°C in 4% BSA. The following day, cells were washed and incubated overnight with cell mask 1: 10 000 (Life Technologies, C10046) and Donkey-anti-rabbit Alexa Fluor 488 (Invitrogen, A21206) 1:10 000 at 4°C overnight in the dark. Cells were mounted using ProLong™ Glass Antifade Mountant with NucBlue™ Stain (Life Technologies, P36981).

Confocal Airyscan images of fixed samples were collected on a Zeiss 880 point-scanning confocal microscope, built on an inverted Zeiss Axio Observer.Z1 stand (Carl Zeiss AG). Images were acquired using a 10x/0.45 Plan-Apochromat objective lens and the Airyscan v1.0 array detector (Carl Zeiss AG).The pinhole for Airyscan imaging was set above 1.25 AU. Multi-channel images were captured sequentially: NucBlue (nuclear marker) using 405 nm excitation and 410-460 nm emission bandwidth, TOMM20 using 488 nm excitation and 500-550 nm emission and CellMask Deep Red using 633 nm excitation and 645-760 nm emission. Images were collected with a 1.8 zoom with an image size of 4300 x 4300 pixels, yielding a pixel size of 0.11µm x0.11µm, and a 1.95µs us pixel dwell time. Images were acquired and processed using the software Zen LSM 2.1 Black (Zeiss), using the Airyscan processing module set to “Auto”.

Images were quantified using ImageJ1.54p. Channels were separated and thresholds were adjusted manually using the “Image/Adjust/Threshold” function. Measurements for cell area, mitochondrial area and nuclear area were assessed through measured setting of cell mask deep red, TOMM20 and NucBlue using the “Analyze/Measure” function on thresholded masks for each channel. Cells which were in the process of dividing or were not captured fully were eliminated from analyses. For generation of mitochondrial area/cell area, individual TOMM20 measured area was divided by the average cell mask area per image.

Representative confocal Airyscan images were collected on a Zeiss 880 point-scanning confocal microscope, built on an inverted Zeiss Axio Observer.Z1 stand (Carl Zeiss AG). Images were acquired using a 40x/1.3 Plan-Apochromat Oil DIC objective lens with Immersol 518 F immersion oil (Carl Zeiss AG), and the Airyscan v1.0 array detector (Carl Zeiss AG). The pinhole for Airyscan imaging was set above 1.25 AU. Multi-channel images of NucBlue, TOMM20 and CellMask Deep Red were captured sequentially as described above. Images were collected with a 1.8 zoom with an image size of 2552 x 2552 pixels, yielding a pixel size of 0.05µm x 0.05µm x 0.22µm, and a 0.76µs us pixel dwell time. Z-stacks were collected using a step size of 5.234 µm and displayed as maximum intensity projections.

Live cell images were stained with 1:1000 Mitotracker green (Thermofisher, M7514). Confocal Airyscan images of live samples were acquired using a 63x/1.3 Plan-Apochromat Oil DIC objective lens with Immersol 518 F immersion oil (Carl Zeiss AG), and the Airyscan v1.0 array detector (Carl Zeiss AG). The pinhole for Airyscan imaging was set above 1.25 AU. Multi-channel images were captured sequentially: MitoTracker Green (mitochondrial marker) using 488 nm excitation and 500-550 nm emission and mCherry using 561 nm excitation and 580-640 nm emission. Representative images were acquired and processed using the software Zen LSM 2.1 Black (Zeiss), using the Airyscan processing module set to “Auto”.

### Labile iron measurement

Cells were seeded at 20.000 cells per well in a sterile black with clear bottom 96 well plate (Greiner, 655090) 4 days after cells were transfected with siRNA. The following day, cells were washed two times with 200 μL HBSS. Just before incubation 1 μmol/L FerroOrange (Insight Biotechnologies, F374-12, Dojindo Laboratories) working solution was prepared in HBSS and 100 μL was added to each well and the plate was incubated for 30 minutes in the 5% CO_2_ at 37C incubator. The fluorescent intensity (Ex. 543 nm, Em. 580nm) was detected using a Tecan plate reader. Afterwards, cells were washed with 200μL PBS, aspirated and plate was put in −80° C. Upon freezing, a protein Bradford assay (Biorad #5000006) was performed using a Tecan plate reader to normalise the fluorescence reading to the protein concentrations of each well.

### Proteomics

Cells were seeded for 60-80% confluence in 10 cm dishes 5 days prior to harvest and transfected with siRNA. Samples were lysed in 100 mM Tris-HCL pH7.5, 4% SDS with 55 mM IAM.

#### Sample preparation for MS analysis

The following methodology was adapted from the procedure described in [65] with only minor modifications. Proteins were reduced with DTT, final concentration 10 mM, to reduce reversible oxidative modifications on cysteine residues, for 1 hour at room temperature, shaking 1,000 rpm, which were subsequently alkylated in the dark with heavy IAA (13C_2_D_2_H_2_INO, Sigma), 55 mM final concentration, 1 hour at room temperature, shaking 1,000 rpm. Alkylated proteins were precipitated with ice cold acetone overnight at −20℃. Precipitated proteins were washed twice with ice cold acetone. Washed pellets were reconstituted in HEPES 200 mM and digested first with Endoproteinase Lys-C (New England Biolabs) for one hour, followed by Trypsin (Promega) overnight. The digested peptides from each experiment, pool or carrier samples, were differentially labelled using Thermo Fisher Scientific TMTpro16 or 18 plex label reagent. 20 µg of each sample was labelled with 0.1 mg of TMT reagent dissolved in 50 μl of 100% anhydrous acetonitrile. The reaction was carried out at room temperature for 2 hours shaking at 1,000 rpm and quenched adding a 5% hydroxylamine solution. Fully labelled samples were mixed in equal amounts and desalted using a 50 mg Sep Pak C18 reverse phase solid-phase extraction cartridge (Waters).

#### High pH peptide fractionation

TMT-labelled peptides were fractionated using high pH reverse phase chromatography on a C18 column (150 × 2.1 mm i.d. - Kinetex EVO (5 μm, 100 Å)) on a HPLC system (Agilent, LC 1260 Infinity II, Agilent). A two-step gradient was applied, from 1–28% B in 42 minutes, then from 28–46% B in 13 minutes to obtain a total of 21 fractions for MS analysis as previously described [66].

#### Nano-UHPLC MS/MS analysis

Peptides in each fraction were further separated by nanoscale C18 reverse-phase liquid chromatography using an EASY-nLC II 1200 (Thermo Fisher Scientific) coupled to an Orbitrap Q-Exactive HF (Thermo Fisher Scientific). Elution was carried out using a binary gradient with buffer A (water) and B (80% acetonitrile), both containing 0.1% formic acid. Samples were loaded with 6 µl of buffer A into a 50 cm fused silica emitter (New Objective) packed in-house with ReproSil-Pur C18-AQ, 1.9 μm resin (Dr Maisch GmbH). Packed emitter was kept at 50 °C by means of a column oven (Sonation) integrated into the nanoelectrospray ion source (Thermo Scientific). Peptides were eluted at a flow rate of 300 nl/min using different gradients which we optimized for three sets of fractions: 1–7, 8–15, and 16–21 as previously described [64] and acquired for 190 minutes. Eluting peptides were electrosprayed into the mass spectrometer using a nanoelectrospray ion source (Thermo Fisher Scientific). An Active Background Ion Reduction Device (ESI Source Solutions) was used to decrease air contaminants signal level. Data acquisition was performed using Xcalibur software version 4.1.31.9 (Thermo Fisher Scientific). A full scan over mass range of 375–1400 m/z was acquired at 120,000 resolution at 200 m/z, with a target value of 3e6 ions for a maximum injection time of 20 ms. Higher energy collisional dissociation fragmentation was performed on the 15 most intense ions selected within an isolation window of 0.8m/z, for a maximum injection time of 96 ms, or a target value of 100,000 ions. Peptide fragments were analysed in the Orbitrap at 45,000 resolution. Former target ions selected for MS/MS were dynamically excluded for 20 seconds.

#### Mass spectrometry raw data analysis

The MS Raw data were processed with MaxQuant [67], *Homo sapiens* (20,675 entries). First and main searches were performed with precursor mass tolerances of 20 ppm and 4.5 ppm, respectively, and MS/MS tolerance of 20 ppm. The minimum peptide length was set to six amino acids and specificity for trypsin cleavage was required, allowing up to two missed cleavage sites. MaxQuant was set to quantify on “Reporter ion MS2”, and TMT16 or 18plex was chosen as the Isobaric label. Interference between TMT channels were corrected by MaxQuant using the correction factors provided by the manufacturer. The “Filter by PIF” option was activated and a “Reporter ion tolerance” of 0.003 Da was used. Modification by Iodoacetamide heavy or light on Cysteine residues (Carbamidomethylation), Methionine oxidation and protein N-terminal acetylation were allowed as variable modifications. No fixed modification was used in the search. The peptide, protein, and site false discovery rate (FDR) was set to 1 %.

#### Quantification and statistical analysis

The text files from the MaxQuant output: “proteinGroups.txt”, “modificationSpecificPeptides.txt”, “peptides.txt” and both heavy and light “Carbamidomethyl (C)Sites.txt” were used for the analysis in Perseus version 1.6.15.0 [68].

The “proteinGroups.txt” file was used for both proteome analysis and for normalization of cysteine containing peptides. The modificationSpecificPeptides.txt file was used as the main table for peptide quantification. The columns with headers “C count”, “Start/End position”, “Length”, “Leading razor protein” from the peptides.txt file, and the columns with headers “Sequence Features”, “Score diff”, “Localisation prob”, “Position” and “FASTA” from the heavy and light Carbamidomethyl (C)Sites.txt files were merged to the modificationSpecificPeptides.txt file using the corresponding id and “Matching by row” tool in Perseus.

The protein or peptides specified in MaxQuant as “Reverse” and “Potential Contaminants” were removed from all tables. For protein quantification, protein groups indicated as “Only identified by site” and those identified with zero unique peptides were removed from the table. Peptides with “Cys count” lower than 1 were excluded from analysis, and only cysteine containing peptides robustly quantified in all the replicate experiments were normalized to the total protein levels and included in the quantification analysis. Only proteins and peptides robustly quantified in all replicates in at least one group, were allowed in the list of quantified proteins. To assess the significance of regulated protein or cysteine peptides between experimental conditions, Student’s t-test or ANOVA were used with a permutation-based FDR below 5%. Welch’s two-sided t-test was performed on log₂-transformed, median-normalised LFQ intensities, with Benjamini–Hochberg FDR adjustment across the proteome. Volcano plots display −log₁₀ nominal p; significance was called as q<0.05 combined with log₂FC > 0.5.

#### OXPHOS classification and assembly-stage assignment

Proteins were classified as core subunits or assembly factors using MitoCarta 3.0 MitoPathways annotations. Each was then assigned to Early, Intermediate or Late assembly stage from canonical biogenesis models — CI: Q-module, Early; Pp/Pd, Intermediate; N, Late [69, 70]; CII: catalytic dimer, Early; membrane anchor, Late [71]; CIII: pre-CIII (cyt-*b* module), Early; UQCRC1/2 core, Intermediate; CYC1/Rieske/UQCR10–11, Late [72]; CIV: COX1/COX2/COX3 modules, Early/Intermediate/Late [73, 74]; CV: F1, Early; peripheral stalk, Intermediate; Fo, Late [75, 76]. Assembly factors were assigned to the stage at which they primarily act. Cytochrome *c* was excluded from the staged analysis. Assembly stage-collapsed log₂FC values were tested against zero by two-sided one-sample Wilcoxon signed-rank test, Holm-corrected across the three stages.

#### Iron-handling protein analysis

A 100-gene panel of iron homeostasis proteins was manually curated covering: IRP/IRE-regulated transcripts (3′UTR and 5′UTR), IRP regulators (IREB2, ACO1, FBXL5), iron transport, mitochondrial iron transport, ferritinophagy (NCOA4), mitochondrial ISC machinery, Fe-S trafficking (CISD1/2/3, BOLA1/2/3, GLRX3), cytosolic Fe-S assembly / CIA machinery, heme synthesis and degradation, heme transport, Fe-S client enzymes (POLA/D/E, PRIM2, BRIP1, DDX10/11, ERCC2, NTHL1, DPYD, PPAT, LIAS, MOCS1, METTL17, CDKAL1, ELP3 etc.) and the NRF2 / glutathione axis (GCLC, GCLM). Full curated list is available (Supplementary Table 3). Differential abundance was derived from Perseus output of LFQ proteomics for ABCB7 vs NTC, with significance defined by Perseus’s permutation-based FDR (S0 = 0.1, FDR = 0.05).

#### Fe-S ligand cysteine accessibility

Per-cysteine fold changes were corrected for protein-level abundance by subtracting the log₂ fold-change of the parent protein from that of the cysteine-containing peptide. Statistical tests were then performed on the corrected values. To remove peptides dominated by Perseus-imputed values (which produce spurious large fold changes), peptides were retained only if at least 3 of 4 replicate values per condition were ≥ log₂ LFQ 11 for both the peptide and the parent protein. A per-replicate residual was calculated for each retained peptide as log₂(peptide LFQ) − log₂(parent-protein LFQ), and Δresidual = mean residual(ABCB7) − mean residual (NTC) was used as the redox-specific effect size. Significance was determined by Welch’s two-sample t-test on per-replicate residuals. Where multiple peptides covered the same cysteine, log₂FCs were averaged and the most significant residual *p*-value retained. Cysteines were classified as Fe-S cluster ligands using UniProt’s metal ion-binding site annotation. Subcellular categorisation (matrix vs non-matrix) was based on MitoCarta 3.0 SubMitoLocalization, with MIM-anchored matrix-facing proteins allocated to matrix group.

### Transcriptional profiling

Quality control of all RNA samples was performed (Agilent Tapestation 4200, High Sensitivity RNA screentape), and only those samples showing RIN values >8 were processed. RNA concentrations were determined by Qubit Fluorometer using the Qubit RNA Broad Range assay (Thermo Fisher Scientific), with 500ng of total RNA used as initial input. Libraries were then prepared using the manufacturers standard procedures (Illumina Stranded mRNA), with Illumina RNA Unique Dual Indexes used to index libraries. Post library QC was then performed using High Sensitivity D1000 screentape (Agilent) for library sizing and profiling and quantified using Qubit High Sensitivity DNA assay. Libraries were then pooled equimolar, to a final concentration of 2nM prior to sequencing. The library pool was then sequenced on an Illumina NextSeq 2000 instrument, at paired end 75bp read length and a depth of approximately 25M reads per sample.

Raw sequencing reads were trimmed using fastp (v.0.23.4) [77] to remove the T overhang. FastQC (v.0.12.1) [56] and MultiQC (v.1.17) [57] were used to assess read quality. Reads were aligned to GRCh38.114 using STAR (v.2.7.11b) single-pass alignment with default parameters [78]. Strandedness was determined using “infer_experiment.py” from the RSeQC package [79]. Gene counts were obtained using featureCounts [80] from the subread package (v.2.0.6) [81]. Differential expression was tested with DESeq2 (v.1.48.1) [82] in R (v.4.5.0) (Wald test, Benjamini–Hochberg FDR < 0.1). Raw counts were normalised using estimateSizeFactors and genes with <10 total reads across all samples were filtered.

#### Gene set enrichment analysis

Pre-ranked GSEA was performed on DESeq2 processed RNAseq data in Python using gseapy.prerank against the MSigDB Hallmark, Reactome and GO Biological Process collections, with genes ranked by the signed −log₁₀(p-value) (sign taken from the log₂ fold-change) and gene sets restricted to 15–500 members; significance was assessed by 1000 gene-set permutations with Benjamini–Hochberg FDR correction. Complete results available in Supplementary Table 4.

### Metabolite profiling

#### Metabolite extraction

Cells transfected or treated as indicated. Four days after, cells were trypsinised, counted and reseeded to reach 80% confluency on day of extraction. The following day, the medium was replenished with fresh medium. After 24 hours steady-state metabolomics and U-^13^C-glutamine labelling was performed as previously published [53]. For U-^13^C-[1]^15^N-cystine-NEM labelling or cystine-NEM metabolomics, medium was prepared using L-cystine-free medium as described, with the addition of either 0.2 μM labelled U-^13^C-[1]^15^N-cystine (Cambridge Isotope Libraries, CNLM-4244-H) or unlabelled L-cystine (Sigma-Aldrich, C8755).

On the day of extraction for steady-state metabolomics and U-^13^C-glutamine, cells were extracted as previously published [52]. For U-^13^C-[1]^15^N-cystine-NEM labelling or cystine-NEM metabolomics, derivatising agent containing 200 μ g/mL N-ethylmaleimide (NEM) (Sigma Aldrich, 04259-5G) and 500ng/ μL ascorbic acid (Sigma Aldrich, A0278-25G) in PBS was added to each well. Samples were mixed at 200 rpm at 4° C for 5 minutes. Samples were washed 3 times with PBS containing 50 μ g/mL NEM with 500 ng/ μ L ascorbic acid. Mixing was at 4° C for 5 minutes for each wash at 200 rpm. Extraction protocol was followed as described elsewhere in methods.

Metabolomics analysis was carried out as previously described [83]. Briefly, chromatographic separation of metabolite extracts was performed using a ZIC-pHILIC column (SeQuant; 150 mm × 2.1 mm, 5 µm; Merck) along with a ZIC-pHILIC guard column (SeQuant; 20 mm × 2.1 mm; Merck) integrated with a Vanquish HPLC system (Thermo Fisher Scientific). A gradient method was utilised, using 20 mM ammonium carbonate (pH 9.2, containing 0.1 % v/v ammonia and 5 µM InfinityLab deactivator (Agilent)) as mobile phase A and 100% acetonitrile as mobile phase B. The elution began with 20% A for two minutes, followed by a linear increase to 80% A over 15 minutes, concluding with a re-equilibration step returning to 20% phase A. The column oven was maintained at 45 °C, with a flow rate to 200 µl per minute. Metabolite analysis and identification were conducted using a Q Exactive Plus Orbitrap mass spectrometer (Thermo Fisher Scientific) equipped with electrospray ionization. The instrument was operated in polarity switching mode at a resolution of 70,000 at 200 m/z, allowing the detection of both positively and negatively charged ions over a mass range of 75 to 1,000 m/z. The automatic gain control (AGC) target was set to 1 × 10^6^, with a maximal injection time (IT) of 250 ms. Data analysis was performed in Skyline (version 23.1.0.455) [84]. Metabolite identification was accomplished by matching accurate mass and retention time of observed peaks to an in-house library generated using metabolite standards (mass tolerance of 5 ppm and retention time tolerance of 0.5 min).

### Mitochondrial immunoprecipitation

siRNA transfected or treated mito-tag U2OS cells were grown in 10 cm dishes. After 5 days, cells were washed twice with ice-cold KPBS and scraped in KPBS (136 mM KCl, 10 mM KH2PO4, pH 7.25, Optima LC/MS grade water). Collected cells were centrifuged at 1,000 x g for 2 minutes at 4°C and supernatant aspirated. Meanwhile, anti-HA Magnetic Beads (Life Technologies) were washed three times with KPBS and put on ice. Each sample pellet was resuspended in 1 mL KPBS, and an input protein sample was collected. The remaining suspension was passed through a 26-gauge needle using 15 strokes. Each sample was centrifuged at 1,000 x g for 3 minutes at 4°C and the supernatant was incubated with 150 μL of washed anti-HA magnetic beads for 3 minutes at 4°C with rotation. Isolated mitochondria bound to magnetic beads were washed three times, using a magnetic stand, with ice-cold KPBS before collecting protein and metabolomic samples.

### Mitochondrial isolation using digitonin

Cells were washed twice in cold PBS and lysed in digitonin buffer (250 mM sucrose, 700 mM Tris-HCl pH 8, 1x Halt protease inhibitors (Life Technologies, 78430) and 100 μg/mL digitonin) for 10 minutes on ice. Lysates were pelleted for 3000 × *g* for 5 minutes to obtain the mitochondrial pellet fraction. The cytosolic supernatant was collected, and the pellet resuspended in RIPA buffer (Life Technologies, 89901) supplemented with cOmplete Mini Tablets and cOmplete Mini Protease Inhibitor Tablets (Roche) and incubated on ice for 15 minutes and centrifuged for 10 minutes at 15,000 rpm. The supernatant was collected as the mitochondrial fraction. Protein concentration in both fractions was determined, and samples were processed as indicated in “immunoblotting”.

### Transfection of mito*GshF* and cyto*GshF*

Bulk populations of empty or SLC25A39KO cells were transfected with pcDNA3 empty plasmid, pMXS-GshF-blasticidin (Addgene, plasmid #210313) or pMXS-mitoGshF-blasticidin (Addgene, plasmid #210314) using Lipofectamine 3000 in 6-wells in triplicate for mtDNA copy number analysis. In parallel, the same cells and plasmids were transfected using 10 cm dishes for western blot analysis. Three days after transfection cells were harvested by trypsinisation and DNA was extracted as described.

### Phylogeny assignments

Each protein was assigned to the deepest taxonomic level at which a *bona fide* ortholog could be recovered using OrthoDB v12.2 [85] (orthologous group IDs and supergroup distributions in Supplementary Table 5). LECA-level placements (ABCB7, SLC25A39, POLRMT, Twinkle, GCN2, AMPKα) were assigned where orthologs were recovered across all major eukaryotic supergroups, including in cases where OrthoDB returned multiple parallel OGs reflecting lineage-specific divergence (Twinkle, GCN2). Opisthokont placements (POLG, TFAM) were assigned where orthology was restricted to animals and fungi, with apparent plant hits attributable to annotation artefacts (POLG) or shared HMG-box domain content rather than functional orthology (TFAM). PGC-1α and PGC-1β were placed at the vertebrate radiation as paralogs arising from duplication of a bilaterian ancestor; family origin was constrained to the bilaterian stem by targeted BLASTP, which recovered a *bona fide* ortholog (both N-terminal coactivator and C-terminal RRM domains; ∼47% identity, E ≤ 1 × 10^−29^) in *Branchiostoma* but only RRM-domain similarity in *Trichoplax* and *Nematostella*, consistent with the previously characterised Spargel ortholog in *Drosophila* [86]. Lineage-age estimates follow standard molecular-clock dates.

### Fly husbandry

*Drosophila melanogaster* samples were cultured in standard medium (1% agar, 1.5% sucrose, 3% glucose, 3.5% dried yeast, 1.5% maize, 1% wheat germ, 1% soybean flour, 3% treacle, 0.5% propionic acid, 0.1% Nipagin). Samples were maintained at 25°C under a 12-hour light/dark cycle. Male flies were used for all experiments unless otherwise stated.

For genetic crosses, 30 virgin females carrying the driver line were crossed with For genetic crosses, 30 adult female virgin flies carrying the driver line were crossed with 15 adult male flies carrying the UAS effector line.

The following UAS effector lines and controls were used:

| Fly line | Source | Description |
| --- | --- | --- |
| w<sup>1118</sup>;UAS-eGFP-IR | VDRC: 60103/KK | RNAi against eGFP. |
| w<sup>1118</sup>;UAS-ABCB7-IR | VDRC: 106039/KK | RNAi against ABCB7. |

Driver lines used for constitutive expression:

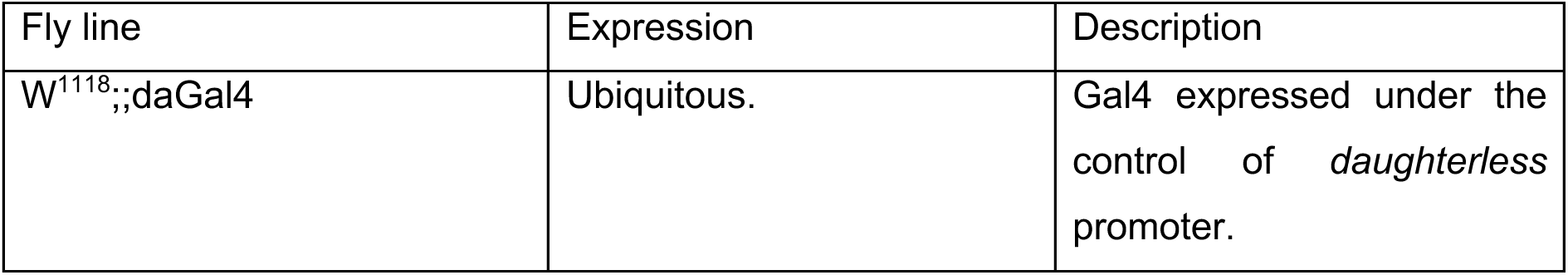

### Arabidopsis growth and DNA extraction

The *Arabidopsis thaliana atm3-1* mutant allele in *AT5G58270* (*ABCB25*) in the Columbia (Col-0) genetic background, which served as wild type, has been described previously [28]. Seeds were surface sterilized, vernalized and grown on agar (0.8% w/v) plates with half-strength Hoagland medium for 15 days in a controlled environment at 21°C with a 16 hour light / 8 hour dark cycle and a photon flux density of 100 - 120 μmol m^-2^ s^-1^. Whole seedlings were harvested and extracted with 200 mM Tris-HCl pH 7.5, 250 mM NaCl_2_, 25 mM EDTA and 0.5% (w/v) sodium dodecyl sulfate (Edward’s solution). DNA was precipitated with 50% (v/v) isopropanol and resuspended in 10 mM Tris-HCl pH 8.0 and 1 mM EDTA.

### Yeast Methods

The *Saccharomyces cerevisiae* (Baker’s yeast) conditional *GAL-ATM1* (*HIS3*) [87] mutant in strain W303-1A (*MATa*, *ura3-1*, *ade2-1*, *trp1-1*, *his3-11*,*15*, *leu2-3112*), used as wild type, were grown on YPGal plates (1% (w/v) yeast extract, 2% (w/v) peptone, 2% (w/v) galactose, 2% (w/v) agar) for 2 days at 30°C. Individual colonies were restreaked and grown on YPD plates (containing 2% (w/v) glucose instead of galactose) to suppress activity of the *GAL1-10* promoter to deplete the Atm1 protein. This was repeated for further depletion, resulting in decreased growth of the *GAL-ATM1* cells. DNA was isolated using the 5-m∼L miniprep method [88].

### Statistics

Samples for metabolomics were randomized by operators. Samples for metabolomics, proteomics and RNA-seq were blinded to the operators. All remaining data collection and analysis were not performed blind to the conditions of the experiments. Data distribution was generally assumed to be normal, with the exception of specific experiments where use of inhibitors or genetic manipulations significantly abrogates measurement, e.g. BSO treatment effect on GSH. In these cases, a non-parametric test (e.g. Kruskal-Wallis) was employed, as described in the figure legends. Data outliers in metabolomics experiments, determined using the Grubbs’ test, were removed before statistical testing. Specific statistical tests used to determine significance, group sizes (*n*) and *p* values are provided in the figure legends. Power calculations were not performed; sample sizes were determined based on field-established norms and are stated in the figure legends. Asterisks indicate significant *p* values in figures, represented as \**p* < 0.05, \*\**p* < 0.01, \*\*\**p* < 0.001 and \*\*\*\**p* < 0.0001. All statistical analyses and data visualisations were carried out using Prism (v.10; GraphPad), RStudio (v.2022.07.0) and Python 3.10, using SciPy (two-sided Welch’s t-tests for two-group comparisons), statsmodels (Benjamini–Hochberg FDR correction), gseapy (pre-ranked GSEA against Hallmark, Reactome and GO-BP), and pandas/NumPy/matplotlib for data handling and plotting; Pearson correlations were used for DepMap co-essentiality and log₂ fold-changes from pre-computed DESeq2 outputs were used for transcriptomic comparisons. Complete figures were then constructed in Adobe Illustrator (v.2026; Adobe).

## Data Availability

Proteomic data are available from PRIDE (PXD078735), metabolomic data are available from MassIVE (MSV000101942).

## Acknowledgements

The authors thank the Core Services and Advanced Technologies at the Cancer Research UK Scotland Institute (grant no. A31287), with particular thanks to Mass Spectrometry (RRID: SCR_027652), Molecular Technologies (RRID:SCR_027368), Flow Cytometry (RRID:SCR_028243) and Beatson Advanced Imaging Resource (RRID:SCR_023875) for their support and assistance with this work. We thank Catherine Winchester for critical reading and editing of the draft manuscript.

## Author contributions

F.M., A.M.S. and P.A.G. conceived the study. F.M. and P.A.G. designed the experiments. F.M. and A.S. performed the CRISPR KO screen and analysis. F.M., A.M.S., J.T.-M. And H.M. performed *in vitro* experiments and analysis. I.F.-G. performed *in vivo* experiments and analysis. P.P. assisted with microscopy experiments and analysis. E.T. and Z.G.G. assisted with *in vivo* characterisation. C.W. performed *in vitro* experiments. M.M. performed *in vitro* experiments and model characterisation. S.L. performed proteomics experiments and analysed the data. A.H.-U. and D.S. performed metabolomics experiments and analysed the data. G.C. performed sequencing experiments and advised on sequencing modalities. J.B. performed *in vitro/in vivo* experiments and supervised analysis. A.S.M. supervised *in vivo* experiments and analysis. P.A.G. supervised the study and obtained funding. F.M. and P.A.G. wrote the paper, with all authors’ involvement and approval.

## Materials

Requests for stable biological materials and sequencing datasets should be addressed to.

---

**Extended Data Figure 1.**
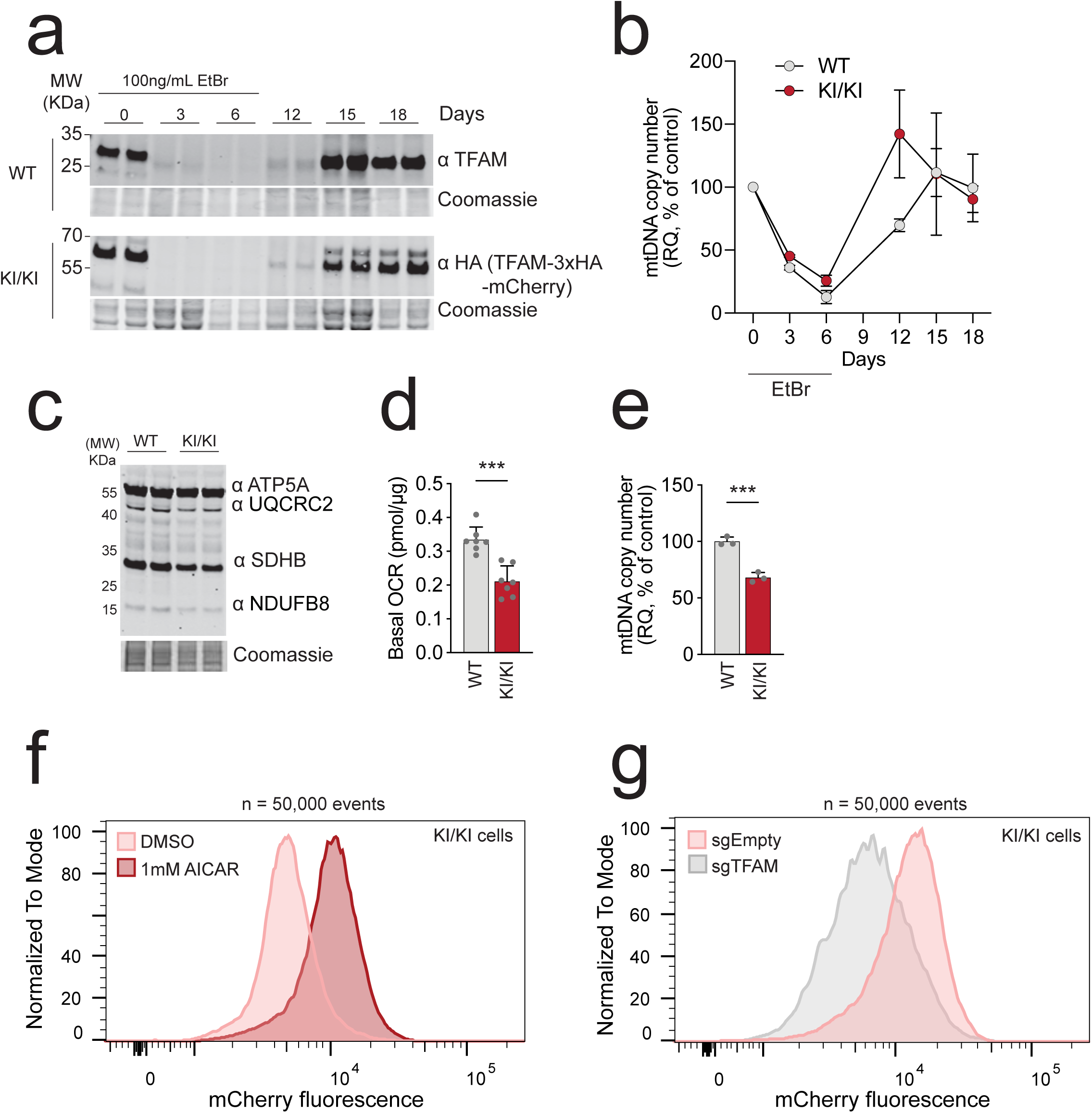
**A.** Immunoblotting of TFAM levels in parental U2OS wildtype and KI/KI TFAM-3xHA-mCherry (KI/KI) cells across ethidium bromide (EtBr) depletion/repletion timecourse. Cells were cultured with EtBr for 6 days to deplete mtDNA and then cultured in medium that did not contain EtBr to allow recovery of mtDNA until day 18. No differences in TFAM depletion/repletion kinetics were observed. Coomassie used as loading control. **B.** Average mtDNA copy number of cells across the time-course, as treated in A. No differences in TFAM depletion/repletion kinetics were observed, indicating that mtDNA transcription and replication were not impacted by the c-terminal mCherry knock-in. Each data point represents the average measurement of three independent wells. Error bars S.D. **C**. Immunoblotting of indicative oxidative phosphorylation (OXPHOS) protein subunits across parental U2OS and KI/KI cells. This experiment was repeated >3 times, a representative result is presented. **D.** Mean basal oxygen consumption rate (OCR) of parental U2OS wildtype and KI/KI cells. KI/KI cells have been clonally selected and demonstrate slightly lower basal respiration. Each data point represents the average of a single well, measured 4 times. Error bars S.D. Welch’s t-test. *** *p* < 0.001. **E**. Mean basal mtDNA copy number of parental U2OS wildtype and KI/KI cells. KI/KI cells have been clonally selected, and demonstrate slightly lower basal mtDNA copy number, in keeping with slightly reduced OCR. Each data point represents the average measurement of three independent wells. Error bars S.D. Welch’s t-test. *** *p* < 0.001. **F**. Flow cytometry histogram of KI/KI cells cultured with 1mM DMSO or 1mM AICAR for 5 days to stimulate mitochondrial biogenesis through AMPK, producing a distinct increase in mCherry fluorescence in the AICAR condition. 50,000 events per group. **G**. Flow cytometry histogram of KI/KI cells transfected with plasmids encoding *S.pyogenes* Cas9 and either an empty sgRNA cassette or sgRNA specific to *TFAM*, demonstrating a distinct decrease in mCherry fluorescence in the sgTFAM condition. 50,000 events per group.

**Extended Data Figure 2.**
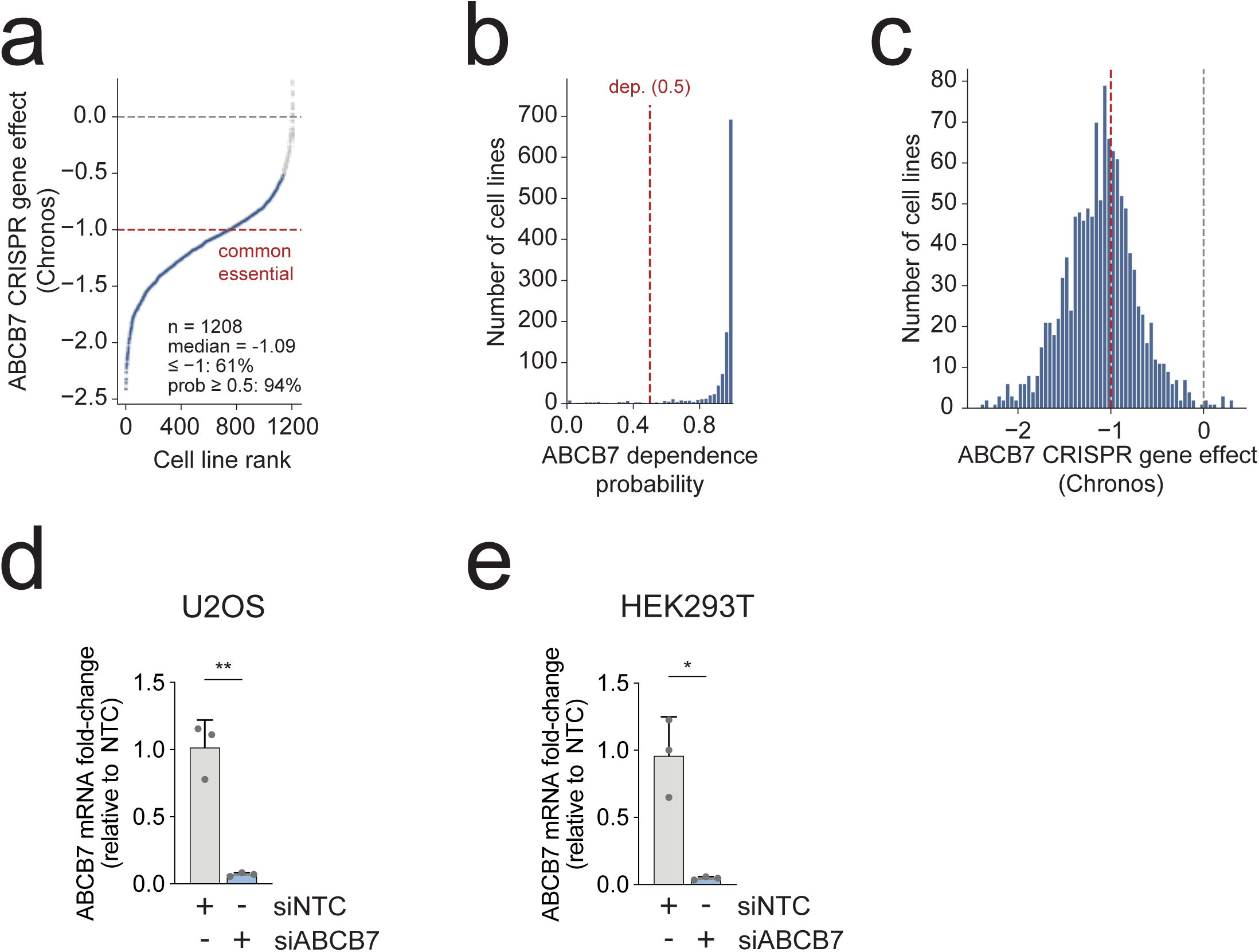
**A.** Waterfall plot of *ABCB7* CRISPR gene effect (Chronos) for n = 1,208 cell lines, ranked from most depleted (left) to least depleted (right); blue, dependency probability ≥ 0.5; grey, < 0.5. Red dashed line, common-essential threshold (−1); grey dashed line, no effect. **B**. Histogram of gene-effect values. Median effect = −1.09; 61% of lines are below the −1 cutoff and 94% are scored as dependent. **C**. Histogram of *ABCB7* Chronos gene effect across n = 1,208 DepMap cell lines (60 bins). Red dashed line, common-essential threshold (−1); grey dashed line, no effect (0). The distribution is unimodal and left-shifted, with a small right tail of non-dependent lines. **D**. Relative fold change of *ABCB7* mRNA in U2OS cells treated with siRNA to *ABCB7* or non-targeting control (NTC) after 5 days. Each data point represents the average of 3 separate wells from each of 3 independent experiments. Error bars S.D. Welch’s t-test. ** *p* < 0.01. **E**. Relative fold change of ABCB7 mRNA in HEK293 cells treated with siRNA to ABCB7 or NTC after 5 days. Each data point represents the average of 3 separate wells from each of 3 independent experiments. Error bars S.D. Welch’s t-test. * *p* < 0.05.

**Extended Data Figure 3.**
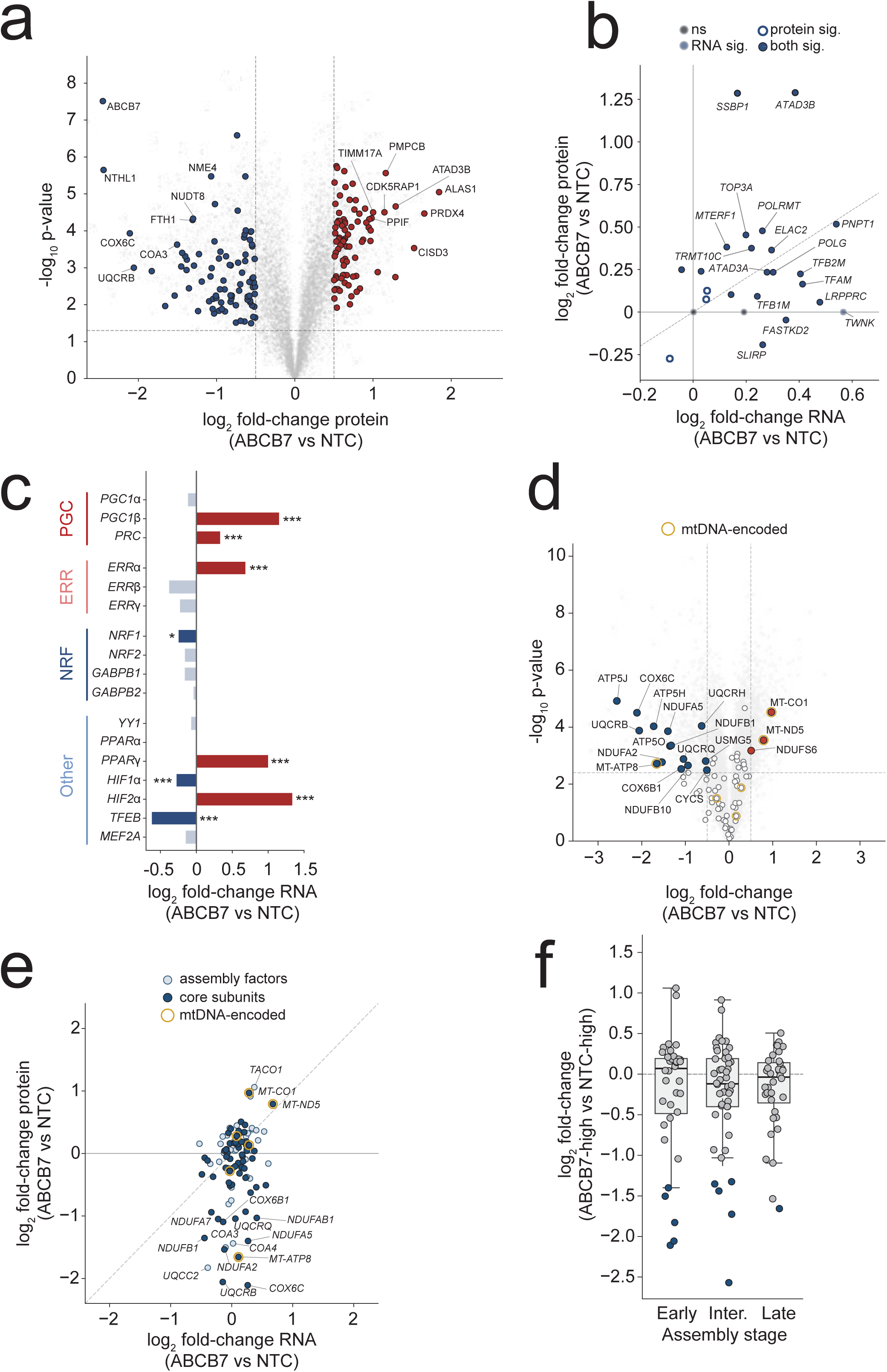
**A.** Volcano plot of all cellular protein abundances. proteins passing Perseus permutation FDR < 0.05 and log₂FC >0.5 are coloured (red, up; blue, down); other proteins in grey. The eight most significantly changed proteins at either end of the distribution are labelled with protein symbols. **B.** Mitochondrial DNA maintenance, transcription and replication machinery scatter plot (RNA vs protein). Filled circles *p* < 0.05 in either layer; open circles n.s. Dashed diagonal: RNA = protein concordance. TWNK is shown on the x-axis (protein not detected). **C.** Canonical mitochondrial biogenesis transcription factor/coactivator RNA abundance. Saturated bars *p* < 0.05; faded bars n.s. Red, up; blue, down. * *p* < 0.05, *** *p* < 0.001. **D.** Differential OXPHOS protein abundance in *ABCB7*-siRNA vs NTC cells. Open circles, MitoCarta 3.0 OXPHOS core subunits (n = 79); filled red/blue, core subunits passing log₂FC > 0.5 and Perseus permutation FDR < 0.05. Gold ring, mtDNA-encoded subunits. **E**. OXPHOS genes (RNA vs protein) scatter plot. The pattern of expression across RNA and protein suggests post-translational differences. Filled dark blue, OXPHOS core subunits; light blue, assembly factors; gold ring, mtDNA-encoded subunits. **F**. OXPHOS-related proteins binned by step in complex assembly (Early/Intermediate/Late; see Methods). Filled blue, proteins passing log₂FC > 0.5 and FDR < 0.05. *p*-values, two-sided one-sample Wilcoxon vs zero, Holm-corrected.

**Extended Data Figure 4.**
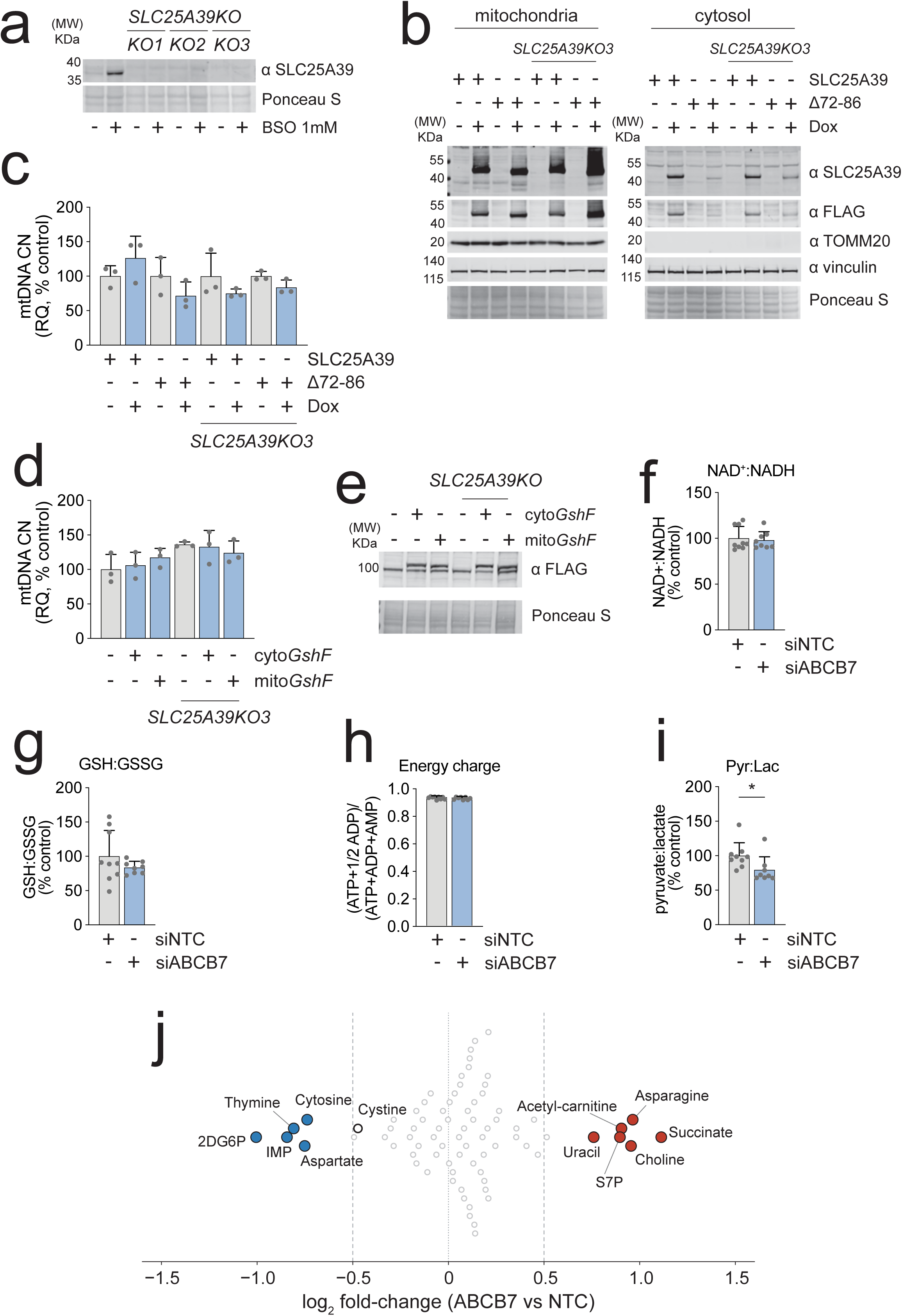
**A.** Immunoblotting of wildtype U2OS and *SLC25A39* knock out (KO) clones for SLC25A39. Cells were treated with 1mM BSO for 48 hours to stabilise SLC25A39 and confirm KO. Ponceau S used as loading control. **B.** Immunoblotting of isolated mitochondria and cytosolic fractions from U2OS wildtype or SLC25A39KO clone 3 cells stably transduced with empty vectors, dox-inducible transgene for *SLC25A39* or a protease-resistant form of SLC25A39, Δ72-86. Cells were induced with 2 μg/mL doxycycline for 24 hours and subjected to digitonin permeabilisation and differential centrifugation. Overexpressed protein (FLAG-tagged) is accumulated in the mitochondrial fraction, as confirmed by TOMM20. Ponceau S and vinculin used as loading controls. **C.** Mean mtDNA copy number analysis of U2OS wildtype and SLC25A39KO clone 3 cells following 5 days of indicated treatments. Each data point represents the mean of 3 wells across 3 independent experiments. Error bars S.D. Two-way ANOVA, Tukey’s multiple comparisons test. **D.** Mean mtDNA copy number analysis of U2OS wildtype and SLC25A39KO clone 3 cells following 5 days of indicated treatments. Each data point represents the mean of 3 wells across 3 independent experiments. Error bars S.D. Two-way ANOVA, Tukey’s multiple comparisons test. **E.** Immunoblotting of U2OS wildtype and SLC25A39KO clone 3 cells transiently transfected with plasmids encoding empty vectors, cyto*GshF* or mito*GshF*. Ponceau S used as loading control. **F.** Ratio of NAD+:NADH from protein normalised LC/MS metabolite profiling. U2OS wildtype cells were treated with siRNA to *ABCB7* or non-targeting control (NTC) for 5 days. Each data point represents a single well across 3 independent experiments. Error bars S.D. **G.** Ratio of reduced to oxidised glutathione (GSH:GSSG) from protein normalised LC/MS metabolite profiling. U2OS wildtype cells were treated with siRNA to *ABCB7* or NTC for 5 days. Each data point represents a single well from each of 3 independent experiments. Error bars S.D. **H.** Energy charge state, calculated from protein normalised AMP, ADP and ATP LC/MS metabolite profiling. U2OS wildtype cells were treated with siRNA to *ABCB7* or NTC for 5 days. Each data point represents a single well from each of 3 independent experiments. Error bars S.D. **I.** Ratio of NAD+:NADH from protein normalised LC/MS metabolite profiling. U2OS wildtype cells were treated with siRNA to *ABCB7* or NTC for 5 days. Each data point represents a single well from each of 3 independent experiments. Error bars S.D. Welch’s t-test. * p <0.05. **J.** Swarm plot of metabolite log₂ fold change in *ABCB7*-KD vs NTC U2OS cells. U2OS wildtype cells were treated with siRNA to *ABCB7* or NTC for 5 days. Per-metabolite log₂FC (*ABCB7* siRNA vs NTC), one point per metabolite (n = 78). Red, significantly up (n = 6); blue, significantly down (n = 5); grey open circles, not significant. Cystine is highlighted (open, black-edged marker) as a metabolite of interest. Significance: *p* < 0.05 (Welch’s t-test on log₂ intensities) and log₂FC > 0.5 (dashed lines). Data points indicated separate wells from each of three independent experiments; one sample excluded from ABCB7 group after Grubbs testing (see Source Data).

**Extended Data Figure 5.**
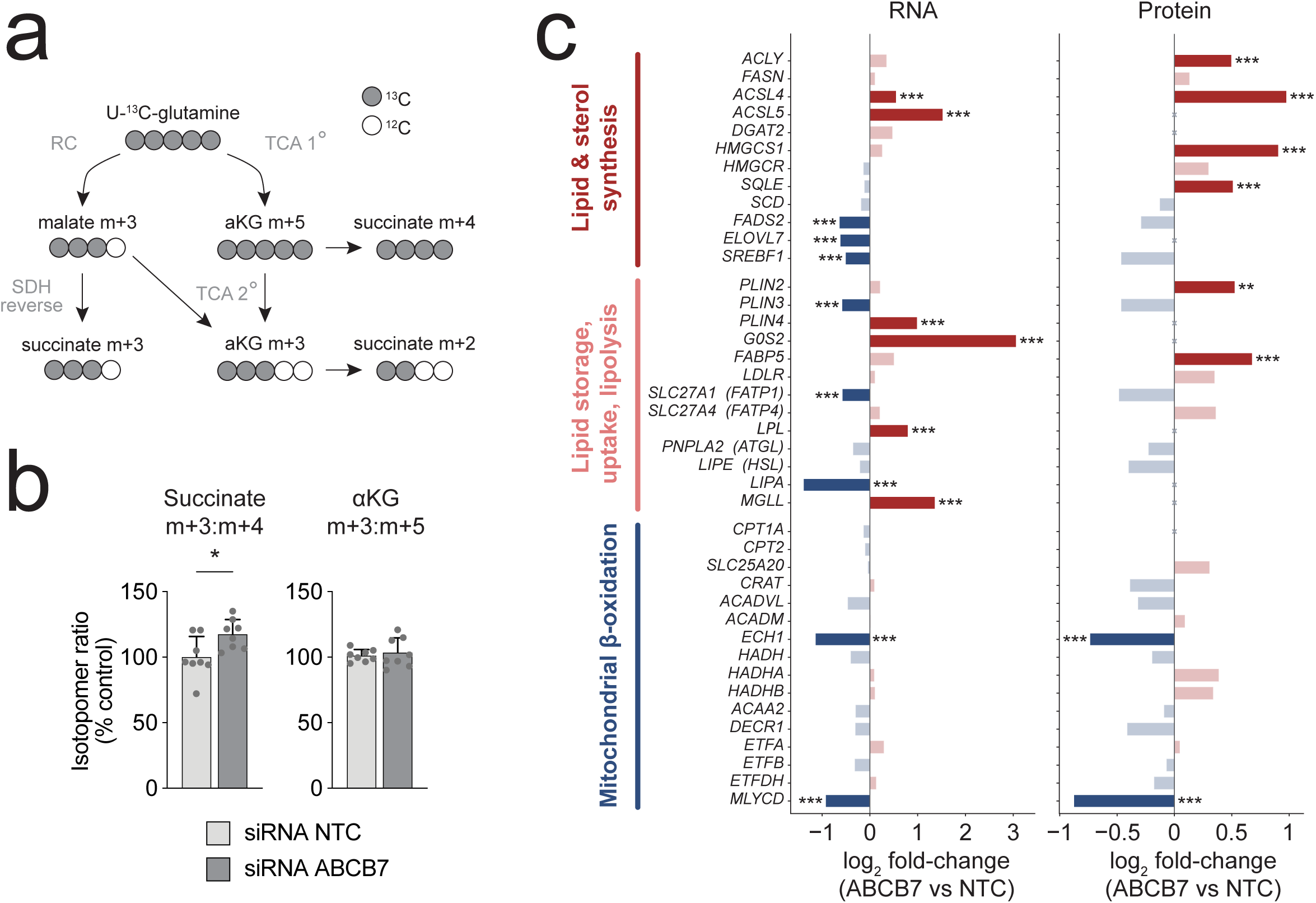
**A.** U-^13^C-glutamine label fate diagram. **B**. Protein normalised ratio of succinate m+3: succinate m+4 isotopomers and aKG m+3:m+5 isotopomers detected by LC/MS. Each data point represents a single well from three independent experiments. Error bars S.D. Welch’s t-test. * *p* <0.05. **C.** Transcript and protein abundances for genes in lipid and sterol synthesis, lipid handling/storage/lipolysis, and mitochondrial β-oxidation after *ABCB7* siRNA. Bar saturation indicates q<0.05 and log₂FC >0.5; faded bars n.s. ✕, not detected. Red, up; blue, down. Coordinate induction of lipid-droplet proteins (PLIN2/4) and the ATGL inhibitor G0S2 with concomitant repression of cytosolic and lysosomal lipases (PNPLA2, LIPE, LIPA) and rate-limiting β-oxidation enzymes (ECH1, MLYCD, ACADVL, DECR1, ETFDH) indicate a shift toward lipid storage and away from oxidation. RNA-level differential expression was assessed with DESeq2 (Wald test) and protein-level differences with two-sided Welch’s *t*-tests on log₂-transformed LFQ intensities; *p*-values were adjusted for multiple testing by the Benjamini–Hochberg procedure within each layer (*q*-values). *p*-value tiers (* *p* < 0.05, ** *p* < 0.01, *** *p* < 0.001) and are placed only on bars passing the *q* < 0.05 threshold. Genes not detected in a given layer are indicated by a grey cross at zero.

**Extended Data Figure 6.**
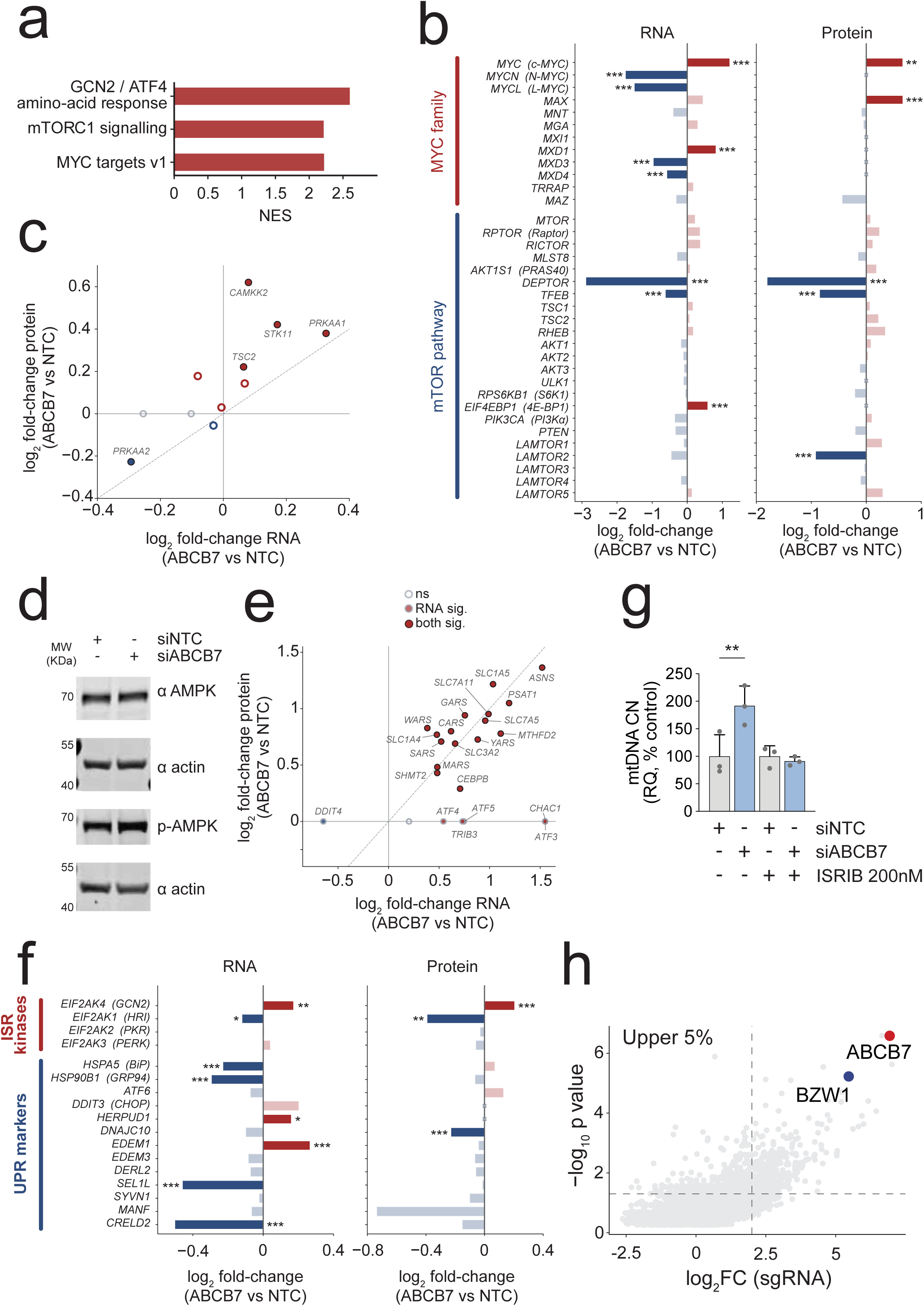
**A.** GSEA against MSigDB Reactome (GCN2/ATF4 amino-acid response) and Hallmark (mTORC1, MYC targets) collections. NES from gseapy, 1,000 permutations. All three pathways FDR<0.001. Full GSEA results available in Supplementary Table 2. **B.** MYC family and mTOR pathway effector transcript and protein abundances. Bar saturation indicates q < 0.05 and log₂ FC > 0.5 (Benjamini-Hochberg FDR; n.s. shown faded). ✕, not detected. Red, up; blue, down. RNA-level differential expression was assessed with DESeq2 (Wald test) and protein-level differences with two-sided Welch’s *t*-tests on log₂-transformed LFQ intensities; *p*-values were adjusted for multiple testing by the Benjamini–Hochberg procedure within each layer (*q*-values) Asterisks denote nominal *p*-value tiers (*** *p* < 0.001, ** *p* < 0.01, * *p* < 0.05) **C.** AMPK subunits, upstream activators (LKB1, CAMKK2) and AMPK-specific substrate (ACACA/B, TSC2, ULK1) scatter plot (RNA vs protein) Filled circles, q < 0.05; open circles, n.s. Red, up; blue, down. Dashed diagonal: RNA = protein concordance**. D.** Immunoblotting of U2OS wildtype cells treated with siRNA to *ABCB7* or non-targeting control (NTC) for 5 days. Actin used as a loading control. This experiment was repeated >3 times, a representative result is presented. **E.** ATF4 / integrated-stress-response marker scatter plot (RNA vs protein): ISR transcription factors (ATF3/4/5, DDIT3, DDIT4, TRIB3, CEBPB), ATF4 amino-acid-program transporters (ASNS, CHAC1, SLC7A11, SLC3A2, SLC7A5, SLC1A4/5), one-carbon metabolism (MTHFD2, PSAT1, SHMT2) and representative cytosolic aminoacyl-tRNA synthetases (CARS, GARS MARS, WARS). Filled circles, *q* < 0.05 in either layer; open circles, n.s. Red, up; blue, down. Dashed diagonal: RNA = protein concordance. **F.** Transcript (left, RNA-seq, DESeq2) and protein (right, LFQ proteomics, Welch’s t-test) log₂ FC for the four ISR effector kinases and canonical UPR markers after *ABCB7* siRNA. Bar saturation indicates *q* < 0.05 and log₂ FC > 0.5 (BH-FDR; n.s. shown faded). ✕, not detected. Red, up; blue, down. Asterisks denote *p*-value tiers (* *p* < 0.05, ** *p* < 0.01, *** *p* < 0.001) and are placed only on bars passing the *q* < 0.05 threshold. Of the four ISR kinases, only EIF2AK4/GCN2 is induced; EIF2AK1/HRI is repressed; PKR and PERK are unchanged. Canonical UPR effectors are not co-ordinately induced, with several constitutive ER chaperones (HSPA5, HSP90B1, SEL1L, CRELD2) repressed at the transcript level. **G.** Mean mtDNA copy number of wild-type U2OS cells treated with siRNA to *ABCB7* or non-targeting control (NTC) after 5 days, supplemented with 200 nM ISRIB. Data points indicate mean mtDNA copy number of three wells across three independent experiments. Error bars, S.D. Two-way ANOVA, Šídák’s corrected. ** *p* < 0.01. **H**. Abundance of sgRNAs in upper 5% TFAM-mCherry fluorescence distribution from initial whole genome CRISPR screen in KI/KI cells (data previously shown in Figure 1g). Thresholds for significance are log_2_ FC < 2, adj-*p* value < 0.05. BZW1, a known competitive antagonist of ISR signalling on the ribosomal pre-initiation complex, is a prominent hit.

**Extended Data Figure 7.**
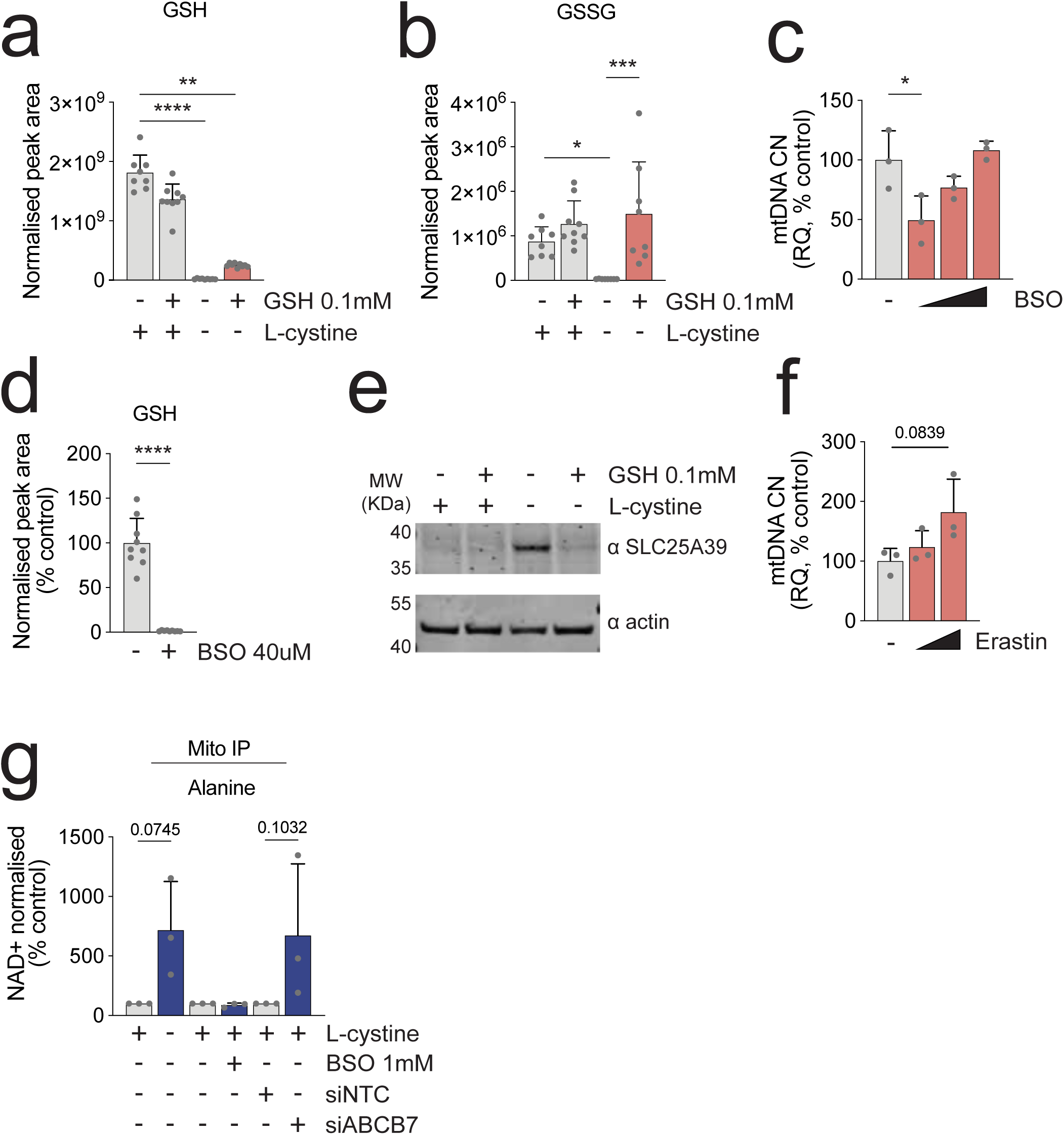
**A.** LC-MS determined mean reduced glutathione (GSH) abundance in U2OS cells grown in cystine replete or depleted medium with or without 0.1mM GSH supplementation for 5 days. Data points indicate three separate wells across three independent experiments. Error bars, S.D. One-way ANOVA, Tukey’s multiple comparisons test *** *p* <0.001, **** *p* < 0.0001. One sample excluded from control group after Grubbs testing (see Source Data). **B.** LC-MS determined mean oxidised glutathione (GSH) abundance in U2OS cells grown in cystine replete or depleted medium with or without 0.1mM GSH supplementation for 5 days. Data points indicate three separate wells across three independent experiments. Error bars, S.D. One-way ANOVA, Kruskal-Wallis test * *p* < 0.05, *** *p* <0.001. **C.** Mean mtDNA copy number of U2OS cells grown in medium supplemented with BSO (sloped triangle indicates 20μM, 40μM, 1mM) for 5 days. Data points indicate mean mtDNA copy number of three separate wells from three independent experiments. Error bars, S.D. One-way ANOVA, Tukey’s multiple comparisons test * *p* < 0.05. **D.** LC-MS determined mean reduced glutathione (GSH) abundance in U2OS cells grown in medium supplemented with BSO for 5 days. Data points indicate three individual wells across three independent experiments. Error bars, S.D. Welch’s t-test. **** *p* <0.0001, **E.** Immunoblotting of U2OS cells grown in cystine replete or depleted medium with or without 0.1mM GSH supplementation for 5 days. SLC25A39 is stabilised by growth in cystine depleted conditions but destabilised by supplementation with GSH. **F.** Mean mtDNA copy number of U2OS cells grown in medium supplemented with Erastin (sloped triange indictes 1μM, 2μM) for 5 days. Data points indicate mean mtDNA copy number of three separate wells from each of three independent experiments. Error bars, S.D. One-way ANOVA, Tukey’s multiple comparisons test **G**. NAD+ normalised mean of alanine abundance from mitochondria purified by mitoIP. Data points are measurements of an individual mitoIP metabolite extraction from three independent experiments. Error bars, S.D. One-way ANOVA, Šídák’s corrected. * *p* < 0.05.

**Extended Data Figure 8.**
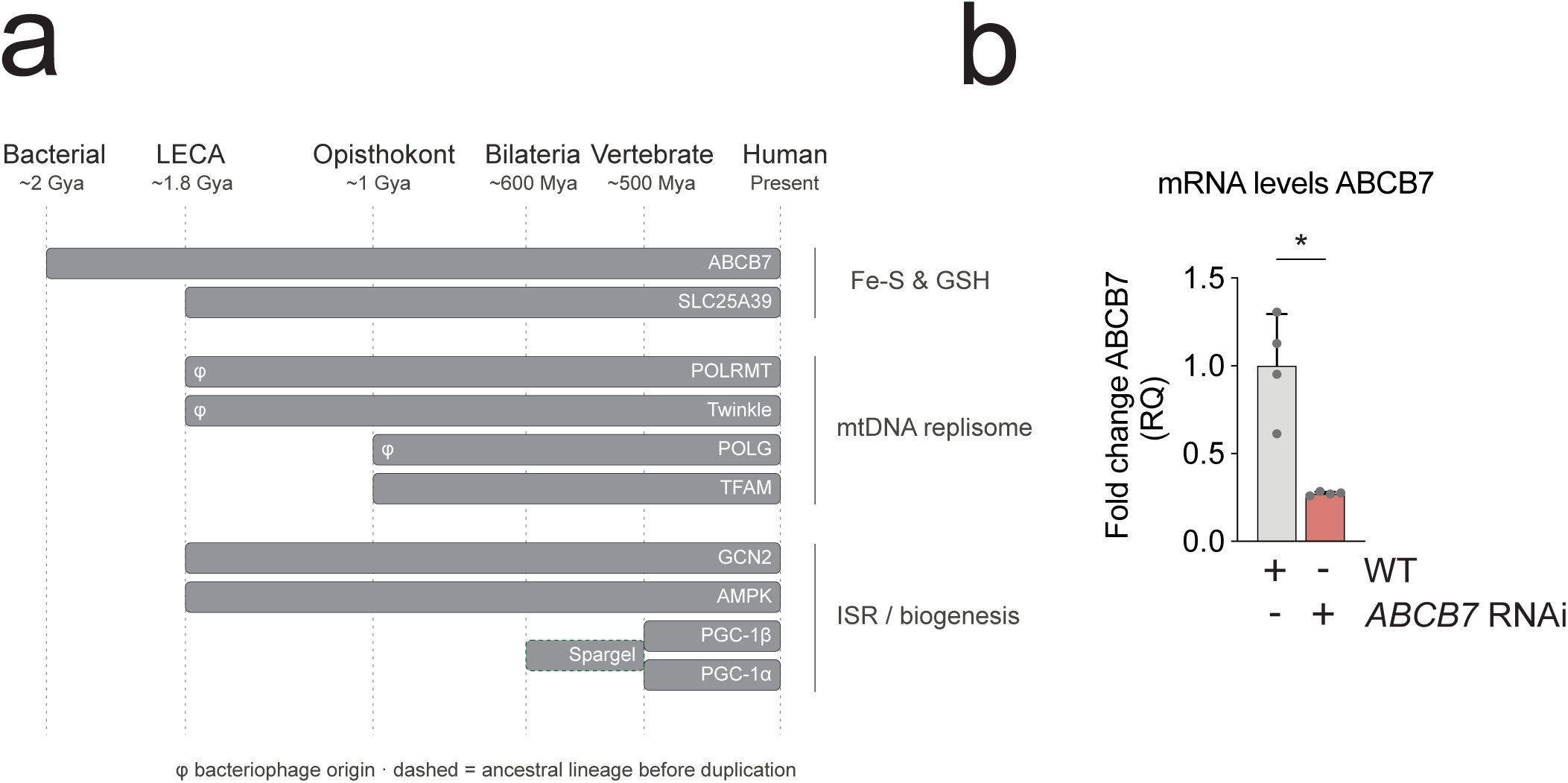
**A.** Evolutionary origins for select elements of the human mtDNA maintenance, mitochondrial stress and biogenesis machinery. Bars span the evolutionary epoch over which each protein’s orthologous group is recovered, from emergence to present day; bar length reflects evolutionary depth. Proteins are grouped by function (Fe-S & GSH, mtDNA replisome, ISR and biogenesis). φ denotes proteins of bacteriophage T7 origin (POLRMT, Twinkle, POLG). The dashed bar indicates the bilaterian PGC-1 family ancestor (Spargel) preceding the vertebrate α/β duplication. ABCB7 alone traces directly to the alphaproteobacterial endosymbiont. Orthology was assigned using OrthoDB v12.2 (Supplementary Table 5) with BLASTP corroboration for SLC25A39 (residue-level conservation of [Fe-S]-coordinating cysteines in the *Arabidopsis thaliana* homolog) and PGC-1α (constraining family origin to the bilaterian stem); see Methods**. B.** Mean fold-change *ABCB7* mRNA level in *D. melanogaster* larvae. Data points indicate individual larvae (n =4). Error bars, S.D. Welch’s *t*-test. * *p* < 0.05.

**Extended Data Figure 9.**
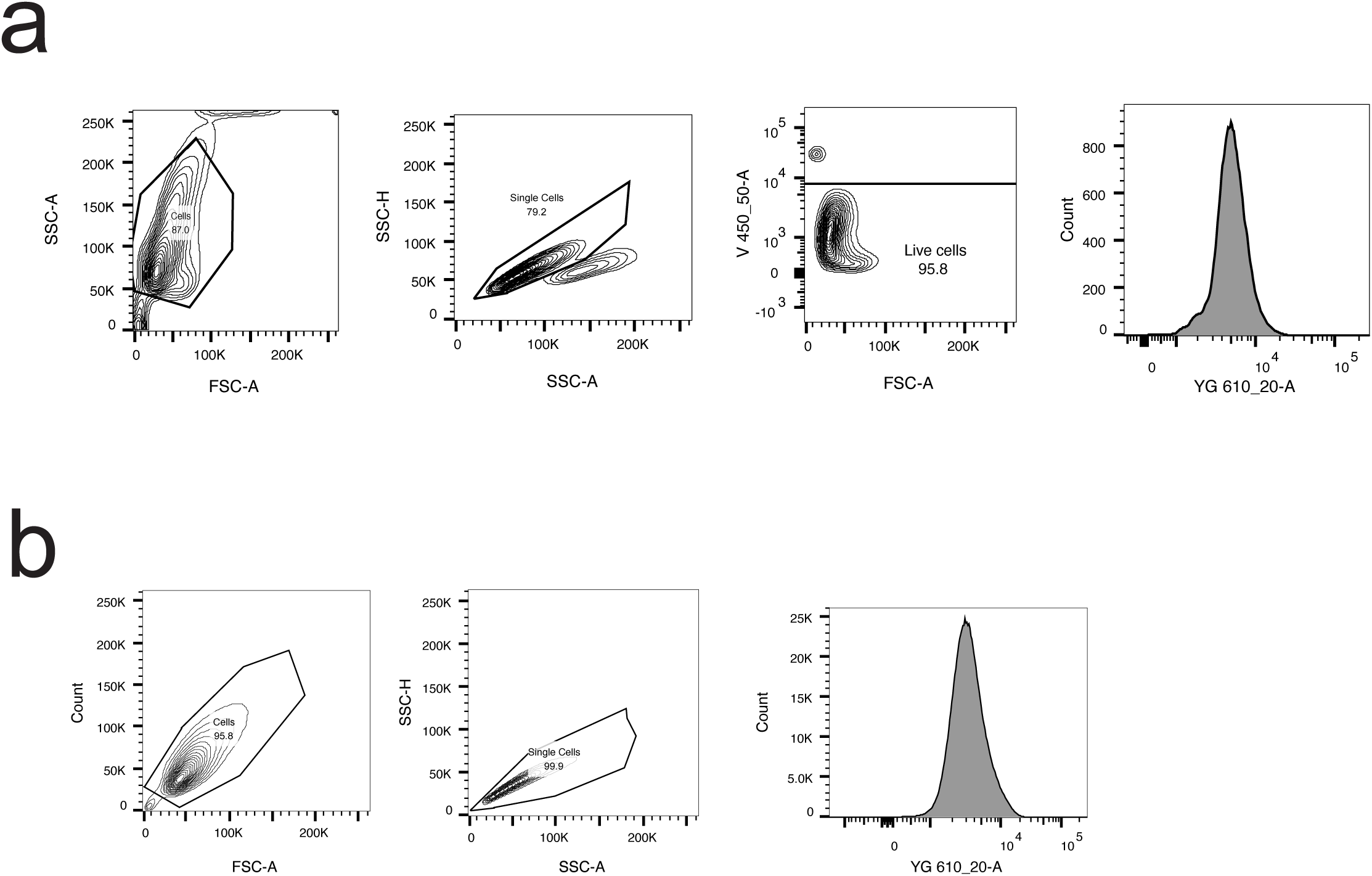
**A.** Flow cytometry gating strategy for live cell analysis presented in Figure 1 and Extended Data Figure 1. Live samples were trypsinised and filtered, before they were stained with DAPI 1:1000. Samples were kept in 4°C in PBS supplemented with 2% FBS before they were run. Live cells: FSC-A vs SSC-A for cells. SSC-A vs SSC-H for single cells. FSC-A vs V450_50-A (DAPI) for live cells. Analysed shift in YG610_20-A (mCherry). Cells were not sorted and were sent to waste. **B.** Flow cytometry gating strategy for fixed cell sorting for CRISPR screen presented in Figure 1. Cells were trypsinised and filtered and then fixed in the dark using a 30-minute incubation, in accordance with the eBioscience™ Intracellular Fixation & Permeabilization Buffer Set (Thermofisher, 00-5523-00). Cells were stored at 4°C in PBS supplemented with 2% FBS. Fixed cells: FSC-A vs SSC-A for cells. FSC-A vs FSC-H for single cells. Sorted cells from YG610_20-A (mCherry).

## References

1. Gupta, R., et al., Nuclear genetic control of mtDNA copy number and heteroplasmy in humans. medRxiv, 2023.

2. Ekstrand, M.I., et al., Mitochondrial transcription factor A regulates mtDNA copy number in mammals. Hum Mol Genet, 2004. 13(9): p. 935–44.

3. Schultz, R.A., et al., Differential expression of mitochondrial DNA replication factors in mammalian tissues. J Biol Chem, 1998. 273(6): p. 3447–51.

4. Nishiyama, S., et al., Over-expression of Tfam improves the mitochondrial disease phenotypes in a mouse model system. Biochem Biophys Res Commun, 2010. 401(1): p. 26–31.

5. Ikeda, M., et al., Overexpression of TFAM or twinkle increases mtDNA copy number and facilitates cardioprotection associated with limited mitochondrial oxidative stress. PLoS One, 2015. 10(3): p. e0119687.

6. Jiang, M., et al., The mitochondrial single-stranded DNA binding protein is essential for initiation of mtDNA replication. Science Advances, 2021. 7(27): p. eabf8631.

7. Fukuoh, A., et al., Screen for mitochondrial DNA copy number maintenance genes reveals essential role for ATP synthase. Mol Syst Biol, 2014. 10(6): p. 734.

8. Burr, S.P., et al., MitoPerturb-Seq identifies gene-specific single-cell responses to mitochondrial DNA depletion and heteroplasmy. Nature Structural & Molecular Biology, 2026. 33(4): p. 711–723.

9. O’Neill, H.M., et al., AMP-activated protein kinase (AMPK) beta1beta2 muscle null mice reveal an essential role for AMPK in maintaining mitochondrial content and glucose uptake during exercise. Proc Natl Acad Sci U S A, 2011. 108(38): p. 16092–7.

10. Arany, Z., et al., Transcriptional coactivator PGC-1a; controls the energy state and contractile function of cardiac muscle. Cell Metabolism, 2005. 1(4): p. 259–271.

11. Puigserver, P., et al., A Cold-Inducible Coactivator of Nuclear Receptors Linked to Adaptive Thermogenesis. Cell, 1998. 92(6): p. 829–839.

12. Wu, Z., et al., Mechanisms Controlling Mitochondrial Biogenesis and Respiration through the Thermogenic Coactivator PGC-1. Cell, 1999. 98(1): p. 115–124.

13. Jäger, S., et al., AMP-activated protein kinase (AMPK) action in skeletal muscle via direct phosphorylation of PGC-1alpha. Proc Natl Acad Sci U S A, 2007. 104(29): p. 12017–22.

14. Scarpulla, R.C., Metabolic control of mitochondrial biogenesis through the PGC-1 family regulatory network. Biochim Biophys Acta, 2011. 1813(7): p. 1269–78.

15. Brüser, C., J. Keller-Findeisen, and S. Jakobs, The TFAM-to-mtDNA ratio defines inner-cellular nucleoid populations with distinct activity levels. Cell Reports, 2021. 37(8): p. 110000.

16. Isaac, R.S., et al., Single-nucleoid architecture reveals heterogeneous packaging of mitochondrial DNA. Nature Structural & Molecular Biology, 2024. 31(3): p. 568–577.

17. Kukat, C. and N.-G. Larsson, mtDNA makes a U-turn for the mitochondrial nucleoid. Trends in Cell Biology, 2013. 23(9): p. 457–463.

18. Ngo, H.B., et al., Distinct structural features of TFAM drive mitochondrial DNA packaging versus transcriptional activation. Nat Commun, 2014. 5: p. 3077.

19. Kang, I., C.T. Chu, and B.A. Kaufman, The mitochondrial transcription factor TFAM in neurodegeneration: emerging evidence and mechanisms. FEBS Letters, 2018. 592(5): p. 793–811.

20. Kukat, C., et al., Super-resolution microscopy reveals that mammalian mitochondrial nucleoids have a uniform size and frequently contain a single copy of mtDNA. Proceedings of the National Academy of Sciences, 2011. 108(33): p. 13534–13539.

21. Alam, T.I., et al., Human mitochondrial DNA is packaged with TFAM. Nucleic Acids Res, 2003. 31(6): p. 1640–5.

22. Filograna, R., et al., Modulation of mtDNA copy number ameliorates the pathological consequences of a heteroplasmic mtDNA mutation in the mouse. Sci Adv, 2019. 5(4): p. eaav9824.

23. Bonekamp, N.A., et al., High levels of TFAM repress mammalian mitochondrial DNA transcription in vivo. Life Science Alliance, 2021. 4(11): p. e202101034.

24. Doench, J.G., et al., Optimized sgRNA design to maximize activity and minimize off-target effects of CRISPR-Cas9. Nat Biotechnol, 2016. 34(2): p. 184–191.

25. Pondarré, C., et al., The mitochondrial ATP-binding cassette transporter Abcb7 is essential in mice and participates in cytosolic iron–sulfur cluster biogenesis. Human Molecular Genetics, 2006. 15(6): p. 953–964.

26. Bekri, S., et al., Human ABC7 transporter: gene structure and mutation causing X-linked sideroblastic anemia with ataxia with disruption of cytosolic iron-sulfur protein maturation. Blood, 2000. 96(9): p. 3256–3264.

27. Schaedler, T.A., et al., A conserved mitochondrial ATP-binding cassette transporter exports glutathione polysulfide for cytosolic metal cofactor assembly. J Biol Chem, 2014. 289(34): p. 23264–74.

28. Bernard, D.G., et al., An allelic mutant series of ATM3 reveals its key role in the biogenesis of cytosolic iron-sulfur proteins in Arabidopsis. Plant Physiol, 2009. 151(2): p. 590–602.

29. Sipos, K., et al., Maturation of Cytosolic Iron-Sulfur Proteins Requires Glutathione*. Journal of Biological Chemistry, 2002. 277(30): p. 26944–26949.

30. Li, J. and J.A. Cowan, Glutathione-coordinated [2Fe-2S] cluster: a viable physiological substrate for mitochondrial ABCB7 transport. Chem Commun (Camb), 2015. 51(12): p. 2253–5.

31. Kispal, G., et al., The ABC transporter Atm1p is required for mitochondrial iron homeostasis. FEBS Letters, 1997. 418(3): p. 346–350.

32. Csere, P., R. Lill, and G. Kispal, Identification of a human mitochondrial ABC transporter, the functional orthologue of yeast Atm1p. FEBS Letters, 1998. 441(2): p. 266–270.

33. Pondarre, C., et al., Abcb7, the gene responsible for X-linked sideroblastic anemia with ataxia, is essential for hematopoiesis. Blood, 2007. 109(8): p. 3567–9.

34. D’Hooghe, M., et al., X-linked sideroblastic anemia and ataxia: A new family with identification of a fourth ABCB7 gene mutation. European Journal of Paediatric Neurology, 2012. 16(6): p. 730–735.

35. Allikmets, R., et al., Mutation of a Putative Mitochondrial Iron Transporter Gene (ABC7) in X-Linked Sideroblastic Anemia and Ataxia (XLSA/A). Human Molecular Genetics, 1999. 8(5): p. 743–749.

36. Maguire, A., et al., X-linked cerebellar ataxia and sideroblastic anaemia associated with a missense mutation in the ABC7 gene predicting V411L. British Journal of Haematology, 2001. 115(4): p. 910–917.

37. Dolatshad, H., et al., Cryptic splicing events in the iron transporter ABCB7 and other key target genes in SF3B1-mutant myelodysplastic syndromes. Leukemia, 2016. 30(12): p. 2322–2331.

38. Casey, J.L., et al., Iron-Responsive Elements: Regulatory RNA Sequences That Control mRNA Levels and Translation. Science, 1988. 240(4854): p. 924–928.

39. Wang, H., et al., FBXL5 Regulates IRP2 Stability in Iron Homeostasis via an Oxygen-Responsive [2Fe2S] Cluster. Mol Cell, 2020. 78(1): p. 31–41.e5.

40. Ben Zichri-David, S., L. Shkuri, and T. Ast, Pulling back the mitochondria’s iron curtain. npj Metabolic Health and Disease, 2025. 3(1): p. 6.

41. Kim, M., et al., Single-cell mtDNA dynamics in tumors is driven by coregulation of nuclear and mitochondrial genomes. Nature Genetics, 2024. 56(5): p. 889–899.

42. Wang, Y., et al., SLC25A39 is necessary for mitochondrial glutathione import in mammalian cells. Nature, 2021. 599(7883): p. 136–140.

43. Liu, Y., et al., Autoregulatory control of mitochondrial glutathione homeostasis. Science, 2023. 382(6672): p. 820–828.

44. Shi, X., et al., Dual regulation of SLC25A39 by AFG3L2 and iron controls mitochondrial glutathione homeostasis. Molecular Cell, 2024. 84(4): p. 802–810.e6.

45. Chen, W.W., E. Freinkman, and D.M. Sabatini, Rapid immunopurification of mitochondria for metabolite profiling and absolute quantification of matrix metabolites. Nat Protoc, 2017. 12(10): p. 2215–2231.

46. Singh, C.R., et al., Human oncoprotein 5MP suppresses general and repeat-associated non-AUG translation via eIF3 by a common mechanism. Cell Rep, 2021. 36(2): p. 109376.

47. Aoyama, K. and T. Nakaki, Impaired glutathione synthesis in neurodegeneration. Int J Mol Sci, 2013. 14(10): p. 21021–44.

48. Ward, N.P., et al., Mitochondrial respiratory function is preserved under cysteine starvation via glutathione catabolism in NSCLC. Nature Communications, 2024. 15(1): p. 4244.

49. Piantadosi, C.A. and H.B. Suliman, Redox regulation of mitochondrial biogenesis. Free Radic Biol Med, 2012. 53(11): p. 2043–53.

50. Couturier, J., et al., The iron-sulfur cluster assembly machineries in plants: current knowledge and open questions. Front Plant Sci, 2013. 4: p. 259.

51. Burger, G., M.W. Gray, and B. Franz Lang, Mitochondrial genomes: anything goes. Trends in Genetics, 2003. 19(12): p. 709–716.

## Method-only Referebces

52. Stemmer, M., et al., CCTop: an agile, open-source network tool for genome-wide prediction of specific CRISPR/Cas9 target sites. 2015: https://cctop.cos.uni-heidelberg.de/.

53. Boscenco, S., et al., Functionally dominant hotspot mutations of mitochondrial ribosomal RNA genes in cancer. Nature Genetics, 2025. 57(11): p. 2705–2714.

54. Herschel, D., Pfeffer, Suzanne R; Jaimon Ebsy. Genomic DNA isolation from fixed cells. 2022 [cited 2024; Available from: https://www.protocols.io/view/genomic-dna-isolation-from-fixed-cells-eq2lynm9qvx9/v1.

55. Marcel, M., Cutadapt removes adapter sequences from high-throughput sequencing reads. EMBnet.journal, 2011. 17.

56. Andrews, S.,. FastQC: A quality control tool for high throughput sequence data. 2010, Babraham Bioinformatics, Babraham Institute, Cambridge, United Kingdom: https://www.bioinformatics.babraham.ac.uk/projects/fastqc/.

57. Ewels, P., et al., MultiQC: summarize analysis results for multiple tools and samples in a single report. Bioinformatics, 2016. 32(19): p. 3047–8.

58. Li, W., et al., MAGeCK enables robust identification of essential genes from genome-scale CRISPR/Cas9 knockout screens. Genome Biol, 2014. 15(12): p. 554.

59. Arafeh, R., et al., The present and future of the Cancer Dependency Map. Nature Reviews Cancer, 2025. 25(1): p. 59–73.

60. DepMap, B., DepMap Public 26Q1.Dataset. 2026.

61. O’Hara, R., et al., Quantitative mitochondrial DNA copy number determination using droplet digital PCR with single-cell resolution. Genome Res, 2019. 29(11): p. 1878–1888.

62. Staneva, D., et al., Yeast Chromatin Mutants Reveal Altered mtDNA Copy Number and Impaired Mitochondrial Membrane Potential. J Fungi (Basel), 2023. 9(3).

63. Ayabe, H., et al., Mitochondrial gene defects in Arabidopsis can broadly affect mitochondrial gene expression through copy number. Plant Physiol, 2023. 191(4): p. 2256–2275.

64. Weber-Lotfi, F., et al., *Mitochondrial DNA Isolation from Plants*, in Mitochondrial DNA: Methods and Protocols, T.J. Nicholls, J.P. Uhler, and M. Falkenberg, Editors. 2023, Springer US: New York, NY. p. 57–75.

65. Lilla, S., et al., SICyLIA-cTMT dissects redox proteome dynamics with high accuracy and depth at microgram scale. Cell Rep Methods, 2025. 5(11): p. 101210.

66. van der Reest, J., et al., Proteome-wide analysis of cysteine oxidation reveals metabolic sensitivity to redox stress. Nat Commun, 2018. 9(1): p. 1581.

67. Cox, J. and M. Mann, MaxQuant enables high peptide identification rates, individualized p.p.b.-range mass accuracies and proteome-wide protein quantification. Nat Biotechnol, 2008. 26(12): p. 1367–72.

68. Tyanova, S., et al., The Perseus computational platform for comprehensive analysis of (prote)omics data. Nat Methods, 2016. 13(9): p. 731–40.

69. Stroud, D.A., et al., Accessory subunits are integral for assembly and function of human mitochondrial complex I. Nature, 2016. 538(7623): p. 123–126.

70. Guerrero-Castillo, S., et al., The Assembly Pathway of Mitochondrial Respiratory Chain Complex I. Cell Metab, 2017. 25(1): p. 128–139.

71. Sun, F., et al., Crystal structure of mitochondrial respiratory membrane protein complex II. Cell, 2005. 121(7): p. 1043–57.

72. Fernandez-Vizarra, E. and M. Zeviani, Mitochondrial complex III Rieske Fe-S protein processing and assembly. Cell Cycle, 2018. 17(6): p. 681–687.

73. Timón-Gómez, A., et al., Mitochondrial cytochrome c oxidase biogenesis: Recent developments. Semin Cell Dev Biol, 2018. 76: p. 163–178.

74. Vidoni, S., et al., MR-1S Interacts with PET100 and PET117 in Module-Based Assembly of Human Cytochrome c Oxidase. Cell Rep, 2017. 18(7): p. 1727–1738.

75. He, J., et al., Assembly of the membrane domain of ATP synthase in human mitochondria. Proc Natl Acad Sci U S A, 2018. 115(12): p. 2988–2993.

76. Walker, J.E., The ATP synthase: the understood, the uncertain and the unknown. Biochem Soc Trans, 2013. 41(1): p. 1–16.

77. Chen, S., et al., *fastp: an ultra-fast all-in-one FASTQ preprocessor*. Bioinformatics, 2018. 34(17): p. i884–i890.

78. Dobin, A., et al., STAR: ultrafast universal RNA-seq aligner. Bioinformatics, 2013. 29(1): p. 15–21.

79. Wang, L., S. Wang, and W. Li, RSeQC: quality control of RNA-seq experiments. Bioinformatics, 2012. 28(16): p. 2184–5.

80. Liao, Y., G.K. Smyth, and W. Shi, featureCounts: an efficient general purpose program for assigning sequence reads to genomic features. Bioinformatics, 2014. 30(7): p. 923–30.

81. Liao, Y., G.K. Smyth, and W. Shi, The Subread aligner: fast, accurate and scalable read mapping by seed-and-vote. Nucleic Acids Res, 2013. 41(10): p. e108.

82. Love, M.I., W. Huber, and S. Anders, Moderated estimation of fold change and dispersion for RNA-seq data with DESeq2. Genome Biol, 2014. 15(12): p. 550.

83. Villar, V.H., et al., Hepatic glutamine synthetase controls N5-methylglutamine in homeostasis and cancer. Nature Chemical Biology, 2023. 19(3): p. 292–300.

84. Adams, K.J., et al., Skyline for Small Molecules: A Unifying Software Package for Quantitative Metabolomics. J Proteome Res, 2020. 19(4): p. 1447–1458.

85. Kuznetsov, D., et al., OrthoDB v11: annotation of orthologs in the widest sampling of organismal diversity. Nucleic Acids Res, 2023. 51(D1): p. D445–d451.

86. Tiefenböck, S.K., et al., The Drosophila PGC-1 homologue Spargel coordinates mitochondrial activity to insulin signalling. Embo j, 2010. 29(1): p. 171–83.

87. Mühlenhoff, U., et al., The yeast frataxin homolog Yfh1p plays a specific role in the maturation of cellular Fe/S proteins. Human Molecular Genetics, 2002. 11(17): p. 2025–2036.

88. Adams, A., et al., Methods in Yeast Genetics. 1997, New York: Cold Harbor Laboratory Press.

